# Uncovering candidate genes involved in photosynthetic capacity using unexplored genetic variation in Spring Wheat

**DOI:** 10.1101/2020.06.15.151928

**Authors:** Ryan Joynson, Gemma Molero, Benedict Coombes, Laura-Jayne Gardiner, Carolina Rivera-Amado, Francisco J Piñera-Chávez, John R Evans, Robert T Furbank, Matthew P Reynolds, Anthony Hall

## Abstract

To feed an ever-increasing population we must leverage advances in genomics and phenotyping to harness the variation in wheat breeding populations for traits like photosynthetic capacity which remains unoptimized. Here we survey a diverse set of wheat germplasm containing elite, introgression and synthetic derivative lines uncovering previously uncharacterised variation. We demonstrate how strategic integration of exotic material alleviates the D genome genetic bottleneck in wheat, increasing SNP rate by 62% largely due to *Ae. tauschii* synthetic wheat donors. Across the panel, 67% of the *Ae. tauschii* donor genome is represented as introgressions in elite backgrounds. We show how observed genetic variation together with hyperspectral reflectance data can be used to identify candidate genes for traits relating to photosynthetic capacity using association analysis. This demonstrates the value of genomic methods in uncovering hidden variation in wheat and how that variation can assist breeding efforts and increase our understanding of complex traits.

## Introduction

Bread wheat occurred through hybridisation of domesticated emmer with diploid goat grass, *Ae. Tauschii* (D)^1^. This event is thought to have occurred very few times in nature, integrating very few *Tauschii* donors and resulting in a genetic bottleneck in D genome diversity^2^. This lack of diversity has been identified in multiple populations, using capture enrichment^3^ and whole genome resequencing^4^ where variation rate in the A/B genomes was >4-fold higher than the D genome. In an attempt to relieve this genetic bottleneck CIMMYT has created >1200 synthetic hexaploid wheat lines^5^ through interspecific crosses of durum wheat *(T. turgidum* ssp. *durum,* AABB) and diverse *Ae tauschii (=Ae. squarrosa)* D genome donors^6^. Many have been used as parents in pre-breeding and breeding programmes, being crossed with elite material producing synthetic-derived lines^7,8^. These lines are attributed to have favourable effects on yield under irrigated conditions^9^, drought stress^10^ heat stress^11^, salinity^12^, biofortification^13^, pre-harvest sprouting resistance^14^ and resistance to several pests and diseases^15^. In addition, introgression of wild relatives has been used to introduce novel diversity with well documented examples such as Rye *(Secale cereale)* and *Thinopyrum ponticum^16,17^.* Despite the broad range of contributions of both synthetic wheat and introgressions to CIMMYTs breeding efforts, little work has been done to characterize the variation in these populations that is hidden to microarray-based techniques that rely on pre-existing knowledge of the variation assayed. Leading to much of this novel genetic variation that has been introduced being overlooked.

In addition to providing diversity for wheat breeders, this genetic diversity can be used to unpick the genetic basis of the traits measured at CIMMYT year on year. We demonstrate this by investigating phenotypic variation in spectral indices that are related to three classes of traits: (i) thermal/hydration properties measured in the infrared part of the electromagnetic spectrum, (ii) pigment related indices assessed in visible bands^18^ and (iii) photosynthesis related indices derived from the whole spectra^19,20^ in the High Biomass Association Panel (HiBAP). Few studies have attempted to determine these traits’ contribution to a plant’s efficiency in utilisation of incident solar radiation (or radiation use efficiency, RUE), which determines crop productivity^21^. Our mechanistic understanding of the genes and pathways involved in RUE is therefore limited, especially under field conditions^22^.

Exploiting existing variation in RUE related traits through identification of the genetic mechanisms responsible could be a straightforward strategy for increasing RUE. A phenotypic range in photosynthetic rates of 33% was observed across 64 winter wheat varieties in UK field conditions^23^ and 50% for 55 spring wheat varieties in Mexico and Australia^24^. To understand the genetic causes of this variation, traits contributing to RUE must be studied. Molero *et al.*, 2019 proposed the use of exotic material (landrace and synthetic-derivative lines) as a resource to increase RUE. Previous work has uncovered multiple marker trait associations (MTAs) related to RUE and biomass accumulation at various phenological stages ^22^ and demonstrated a link between RUE and photoprotection.

This suggests that photo-protective pigments could contribute to RUE throughout the crop cycle by preventing the propagation of free radicals that damage photosynthetic machinery. In addition, photosynthetic potential may differ depending on the content of individual leaf pigments^25^ as the amount of solar radiation absorbed depends on pigment content^26^ which in turn relates to photosynthetic capacity^27^. Chlorophyll *a* (chl*a*) is the primary pigment of photosynthesis while chlorophyll *b* (chl*b*) is an accessory pigment. In a study of Australian wheat varieties released through time, a decrease in the Chl a/b ratio was associated with a decrease in electron transport capacity per unit of chlorophyll, but because total Chl content per unit leaf area increased, electron transport capacity per unit leaf area increased^28^. Assessment of the contribution to RUE from pigment composition and its underpinning genetic basis is, therefore, of great interest for enhancing photosynthetic potential of wheat.

Through enrichment capture and *de novo* SNP discovery we are able to gain an unprecedented insight into the overall levels of genetic diversity within CIMMYT breeding material. Using this data, we can determine the contribution of *Ae. Tauschii, S. secale* and *T. ponticum* donors to increasing diversity in exotic derived lines. We have utilised this novel genetic information to further investigate the link between photoprotection and RUE through genome wide association of leaf pigment compositions of 149 wheat lines using high throughput hyperspectral reflectance measurements taken over 2 growing seasons. This uncovered novel MTAs for >20 traits relating to leaf pigmentation and water content along with candidate genes containing possible causative non-synonymous variants that could be leveraged for improvements in RUE.

## Results

### Genotyping and SNP effects

To investigate genetic variation across the HiBAP panel we used a *de novo* SNP discovery strategy using a bespoke target sequence capture design. We developed a 12-Mb target sequence using the MyBaits system based on that described by Gardiner et al., 2018^3^, where underperforming baits were replaced with baits targeting genes associated with photosynthesis and biomass accumulation. A schematic of this capture design can be seen in **SF1**.

In total, 18.6 billion reads were sequenced between the 149 lines, an average of 124 million per line. Of these, 86% mapped uniquely covering 420Mbp of the genome to 5x or greater and 172Mbp at 10x or greater. A breakdown of mapping efficiency and variant calling can be found in **ST1**. Variant calling yielded an average of 764,825 homozygous SNPs per line, producing a marker density of 45 SNPs/Mb. Of these, 96.9% of SNPs were in intergenic regions and 3.1% in the genic regions. Of the SNPs in gene bodies, 49% resulted in synonymous substitutions and 51% in non-synonymous substitutions (**ST2**). The average number of SNPs for the panel members containing exotic pedigree history was 11% higher than that of the elite subpopulation overall. The exotic population showed an increased SNP rate in all subgenomes, 5%, 10% and 62% for the A, B and D genomes respectively. A T-test comparing the elite and exotic subpopulation showed no significant differences between the number of reads nor the number of bp that were mapped to ≥5x coverage between populations. After filtering for SNP loci with <10% missing data for MAF of >5%, 241,907 shared loci were retained. Overall marker density across the shared variants was 17 SNPs/Mbp with the highest density in the B genome followed by the A and D with 25, 16 and 9 SNPs/Mbp respectively **(ST3)**. Genome-wide SNP subset density can be seen in **Figure 1** and **SF2**.

**Figure 1.**
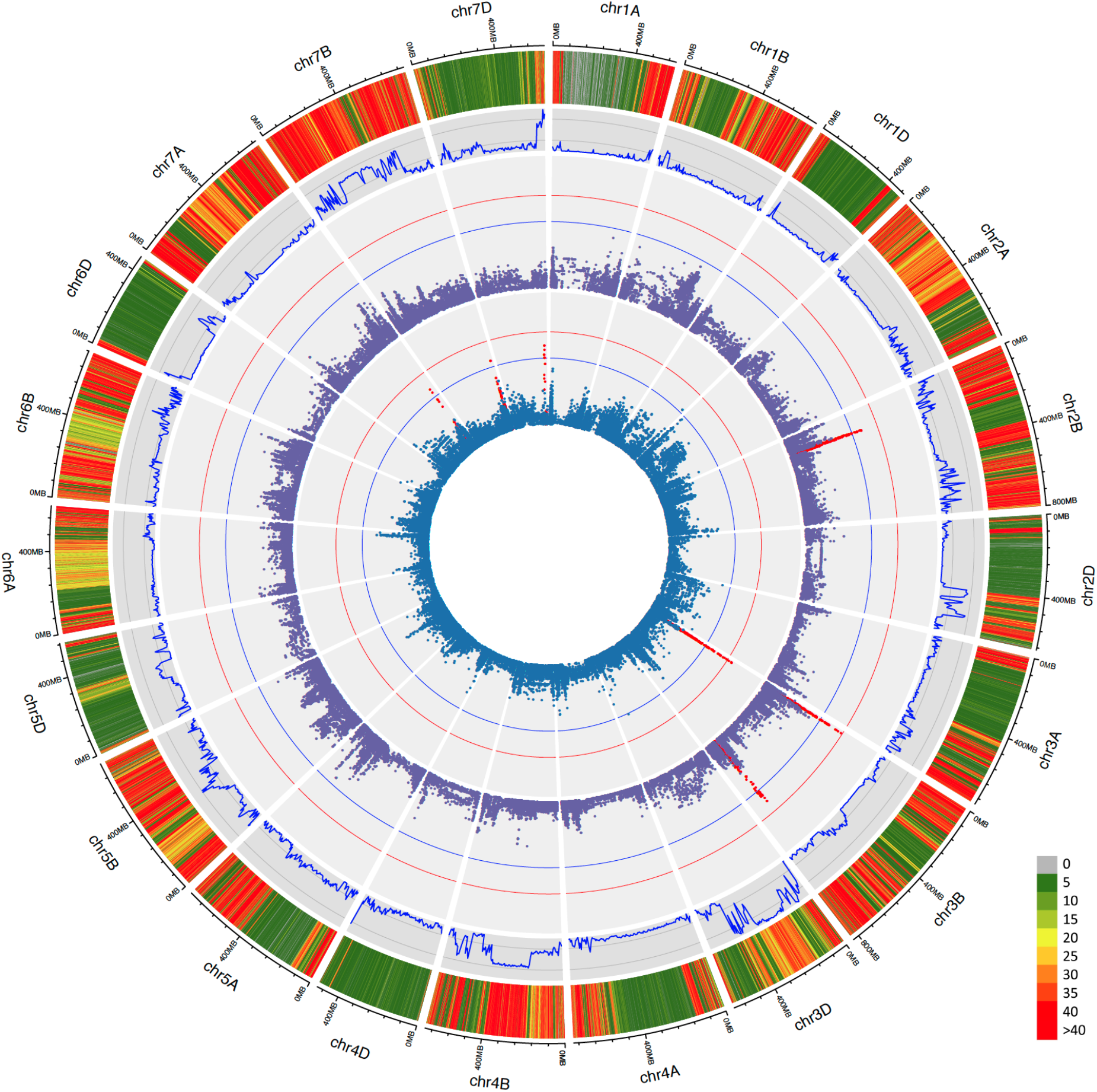
Genetic analysis of the HiBAP panel. (From outside to inside) A) SNP density heatmap across the genome of loci containing <10% missing data and >5% MAF within the HiBAP panel in 100Kbp bins. B) Fixation index calculated between elite background and exotic background subpopulations. C) Genome wide association of flag leaf chlorophyll b content (C) Genome wide association of carotenoid content. Significance cut-offs for -log10p of 5 and FDR correction are shown as blue and red lines respectively. SNPs in an interval above significance thresholds are shown in red.

### Population structure analysis

Model-based Bayesian clustering methods were used to deduce the population structure of the panel. The Evanno method revealed evidence for 2 subpopulations and some evidence for as many as 8 subpopulations (**SF3**). Where 2 subpopulations are assumed, population 1 and 2 comprise 114 and 35 respectively (**SF4**). Of population 2 members, 88.5% had synthetic/landrace parents in their pedigree history whereas only 16% of population 1 had any exotic background (**Figure 2**). Multiple lines also demonstrated significant admixture between populations. Admixture was seen to a lesser extent in the elite backgrounds (4%) compared with exotic backgrounds (25%). Fst analysis demonstrated genomewide effects of integration of exotic material with large regions of chromosomes showing differences between the elite background and exotic background panel members. Most notably in chromosomes 2D, 3D, 4B and 7B with regions spanning >300Mbp (**Figure 1**/**SF5-A**). When the elite population was split randomly into two pseudo-populations, Fst was negligibly small across every chromosome (**SF5-B**).

**Figure 2.**
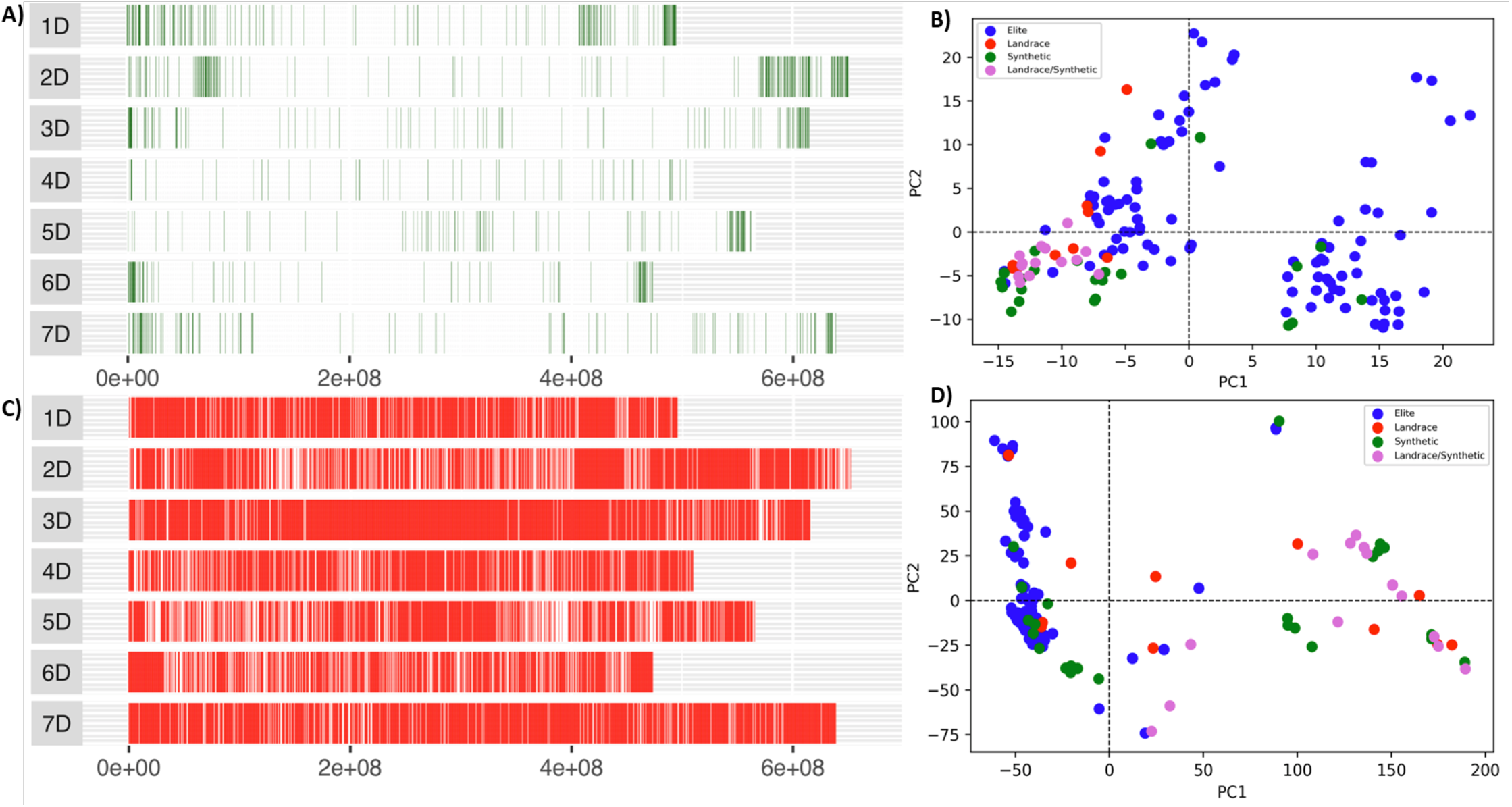
Enrichment capture reveals hidden variation contributed by exotic material. Distribution of D genome polymorphic SNP markers in the HiBAP panel from A) The 35K wheat breeders’ array and B) PCA demonstrating the identified genetic variation using the 35K array SNPs C) *De novo* SNP distribution from enrichment capture data after filtering the combined panel data for < 10% missing data and a minor allele frequency (MAF) of > 5%. D) PCA demonstrating the identified genetic variation using the *de novo* called enrichment capture genotyping SNPs

### Synthetic wheat introduces substantial increase in D genome variation

Comparison of SNP density in the D genome within the elite and exotic subpopulations revealed an increase of 62% in variation in the exotic subpopulation **(Table 1)**. The largest increase was seen on chromosome 3D, with an increase of ~200%. However, these increases were not universal, with 1D showing no notable increase in SNP numbers between populations. Comparison of the elite and exotic subpopulation members highlights that these increases in D genome SNP density is localised into blocks (**Figure 3**), a result of the low crossover rate of 1-2 per chromosome per cross in wheat ^29^, with regions as large as 343Mb showing notable SNP density increases. Within these regions a large proportion of the SNPs matched variation called for modern *Ae. Tauschii,* confirming the origin of this variation is the *Ae. Tauschii* donor used in synthetic creation (**Figure 3B**). The overall size of donor regions varied widely across the synthetic subpopulation, spanning from 43% to as little as 0.5% of the D subgenome and differed from the theoretical D contribution to pedigree range (1.6 – 25%) estimated from a dilution factor associated with the number of crosses after the original cross with the primary synthetic (**ST4**). In 13 members with synthetic backgrounds negligible levels of donor were identified (0.5-5%), suggesting donor loss during subsequent crosses and selections for agronomic traits/ideotypes. This loss was not necessarily correlated with the theoretical D pedigree contribution. For example, lines HiBAP_49 and HiBAP_51 are sister lines derived from the same cross (SOKOLL//PUB94.15.1.12/WBLL1) with a common selection history and a theoretical pedigree contribution of 6.3% from *Ae. Tauschii,* but they contain 12.7% and 26.0%, respectively of *Ae. Tauschii* regions in their D subgenome (**SF6 and ST4**). Elite members showed almost no regions of increased diversity to the extent seen in those with synthetic history when compared to the CS reference (**Figure 3A**). All D subgenome chromosomes are seen to contain regions of greatly increased SNP density in at least one member of the synthetic history subpopulation. Across all panel members with synthetic pedigree history 5301 non-redundant 500Kbp bins were identified with a 5-fold increase in variation when compared to the average number of SNPs for each bin across the elite subpopulation. This equates to ~2.65Gbp (67.1%) of D genome sequence within the synthetic subpopulation that are likely to originate from donor *Ae. Tauschii.* These regions encompass 22,583 high confidence genes in the CS reference annotation.

**Table 1.**
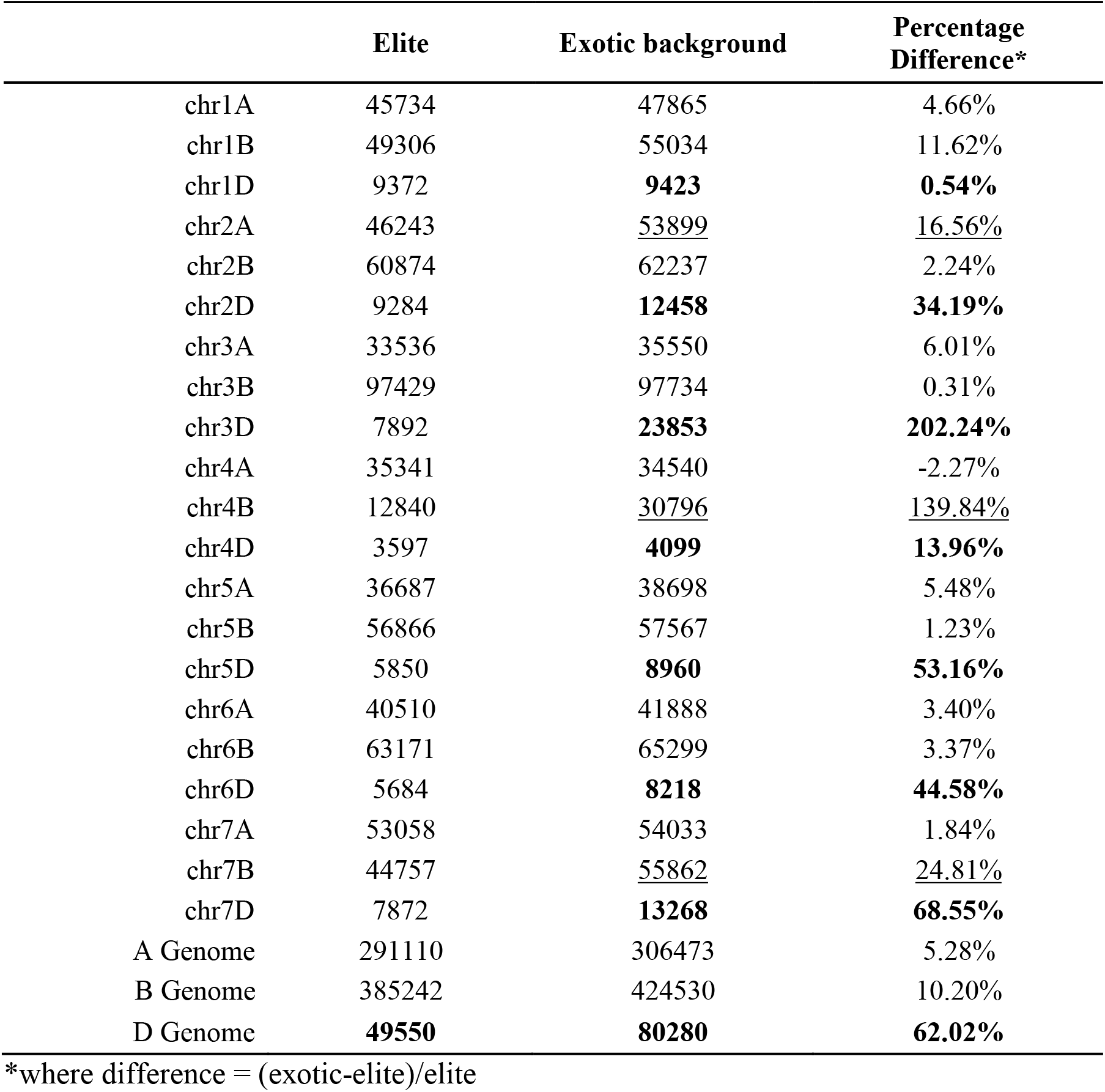
The average number of de novo SNP calls per chromosome for the elite and exotic subpopulations

### Wild relative introgressions can be tracked using de novo SNP calls

To identify introgressions from Rye *(Secale cereale),* SNPs from each line were separated into 500kbp bins for all subgenomes and the number of variants that match Rye in both position and allele were counted. SNPs between Rye and Chinese spring (CS) were generated by mapping and variant calling Lo7 Rye Illumina sequencing reads (ERS446995) against the CS reference genome. This revealed the 1B/1Rs introgressions in 6 panel members spanning the first 239Mb in each line (**SF7**). This region contains 1507 high confidence genes in the CS reference genome annotation (v1.1). An introgression on the long arm of chromosome 7D was also identified in 3 panel members that spans a 300Mb region from 344Mb to the end of the chromosome (**SF8)**; this interval contains 2563 genes in the CS reference annotation. The pedigree history of these lines suggests this is an introgression from *Thinopyrum ponticum*. Panel members containing *S. Secale*/*T. ponticum* introgressions can be seen in **ST5**.

### Phenotypic variation for N content, and spectral indices

The results from analysis of variance (ANOVA) for N content, vegetation indices, pigment composition, senescence, water indices and traits estimated from wheat physiology predictor indicated significant variation among genotypes, environments, and genotype × environment interactions with few exceptions (**Table 2**). Broad sense heritability (*H*^2^) was high for NLamA7, medium for SPAD_A7_, low to medium for vegetation indices, pigment composition and senescence/degradation indices, high to medium for water indices and low for LMA and RDM (**Table 2**). Broad phenotypic variation among genotypes in spectral reflectance between the visible spectrum and the distribution of values for carotenoid, *chla/b* content was observed (**Figure 4**).

**Figure 3.**
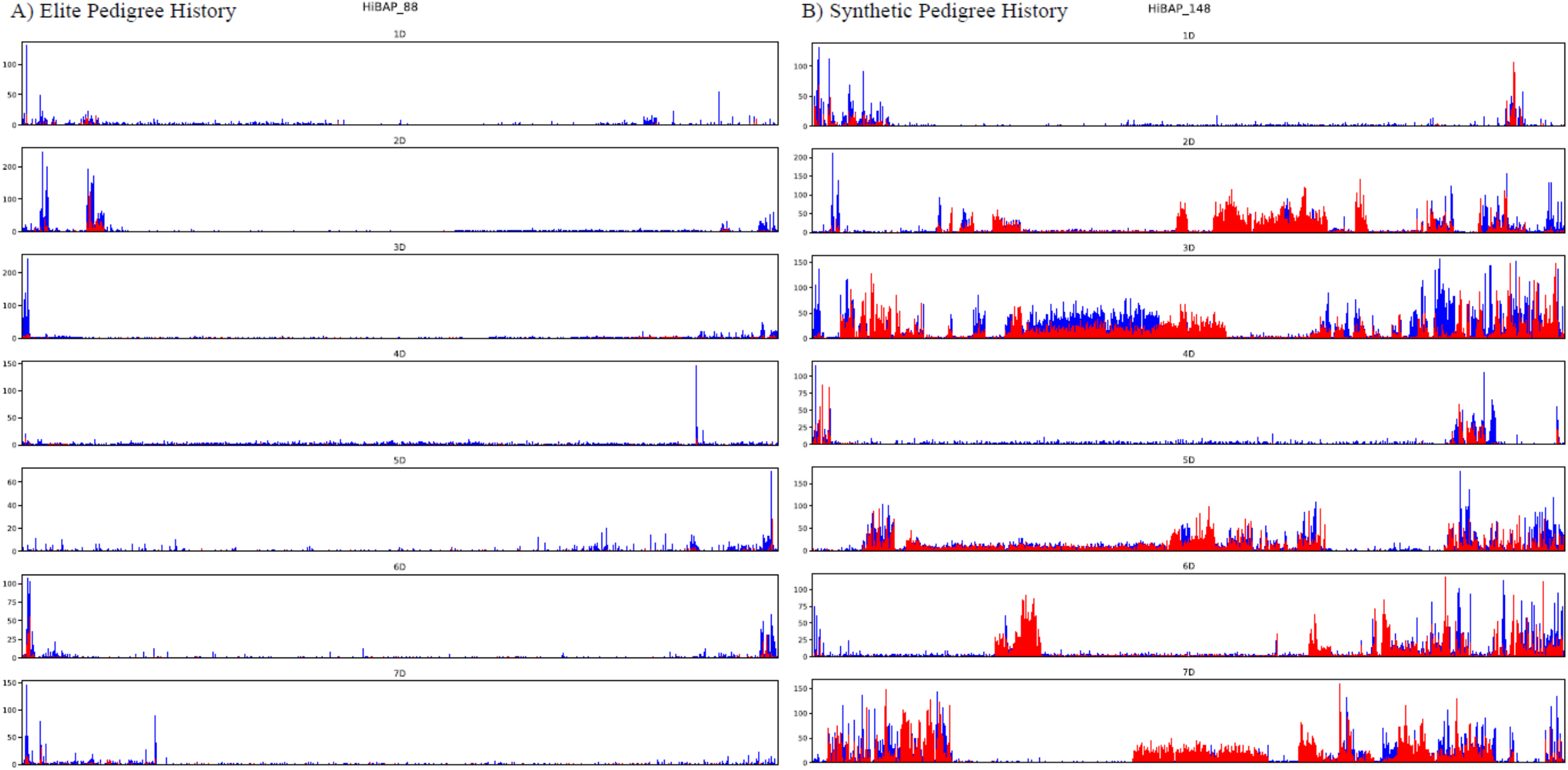
Synthetic wheat donor introgression identification in the D genome. D genome SNP density plots for A) a representative example of HiBAP elite population and B) an example of a member of the synthetically derived subpopulation. SNPs were binned into 500Kbp bins, demonstrating the number of SNPs that matched in position and allele of those seen in *Ae. Tauschii* against Chinese Spring reference genome (red) and the number of SNPs that did not match (blue).

**Figure 4.**
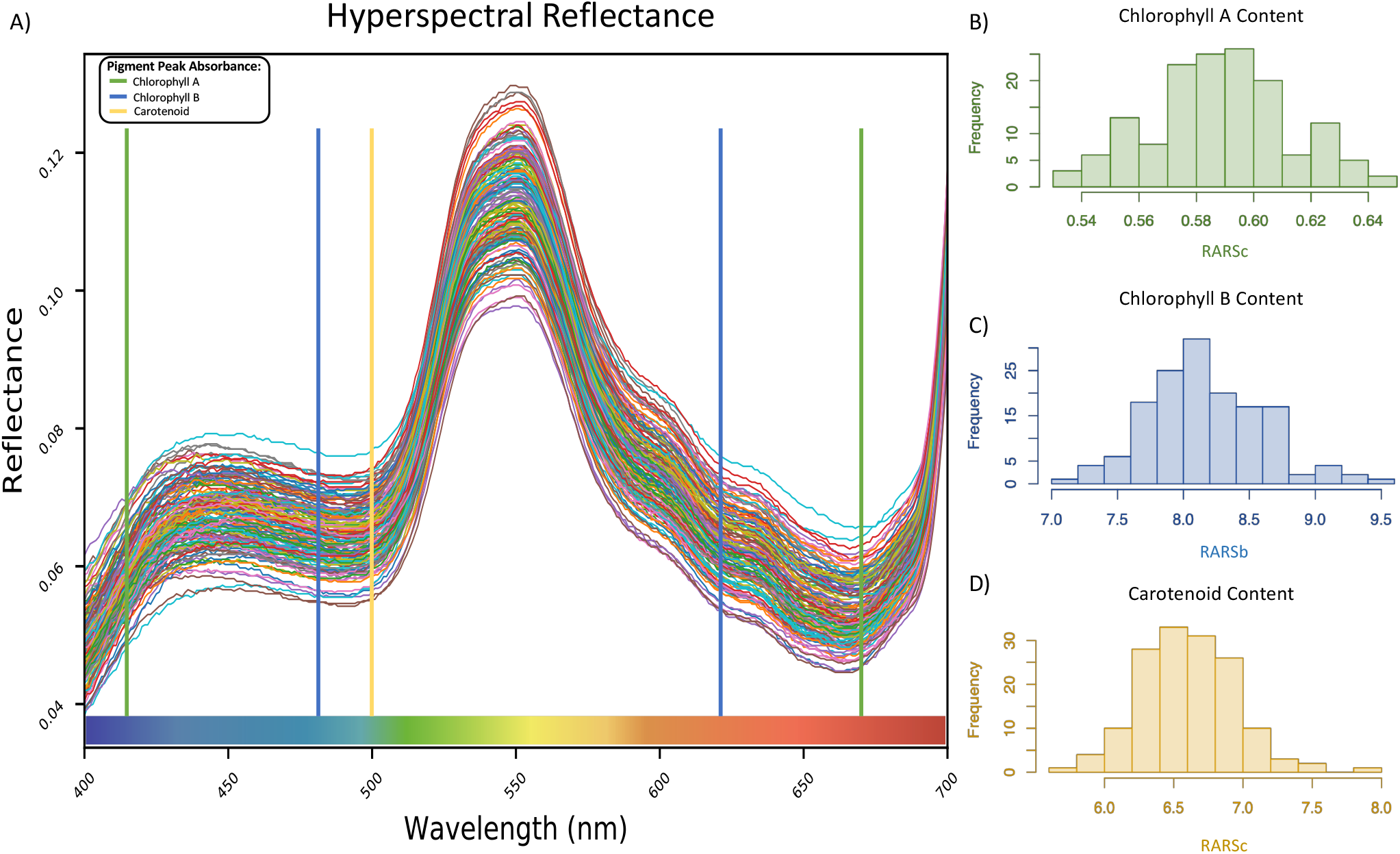
Phenotypic variation in Spectral reflectance from the 149 lines: A) The level of variation in reflectance of the visible portion of the hyperspectral reflectance data for each member of the HiBAP panel. The distribution of observed values derived from spectral indices B) flag leaf chlorophyll a C) flag leaf chlorophyll b and D) flag leaf carotenoid content across all panel members where frequency relates to the number of panel members within each bin in each histogram.

**Table 2.**
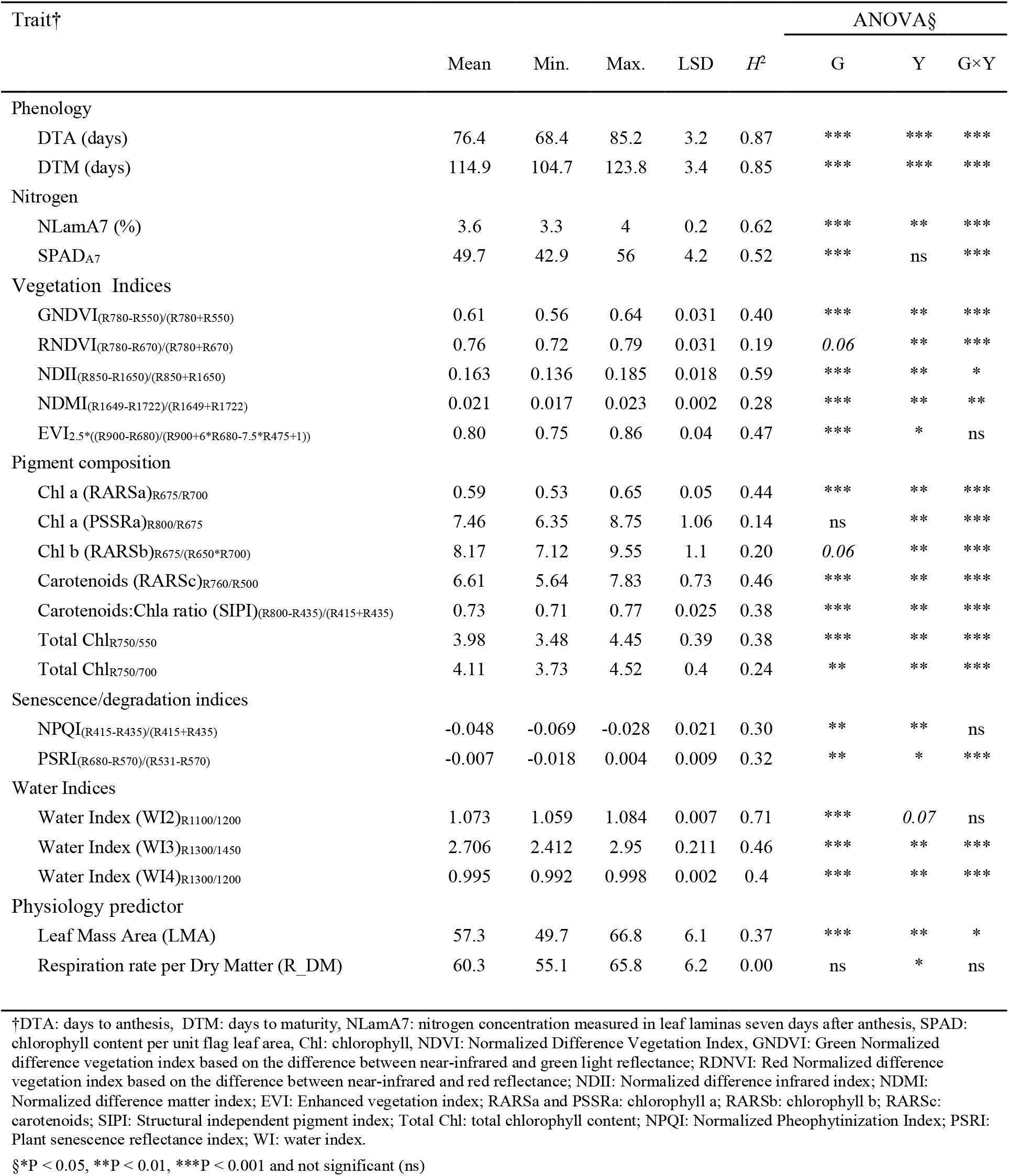
Descriptive statistics, broad sense heritability (*H*^2^) and ANOVA for phenology, nitrogen content and hyperspectral indices of HiBAP grown for two years (Y15-16 and Y16-17) in northwest Mexico under full irrigated conditions.

### Association between spectral indices and agronomic traits

Multiple regression analysis (stepwise) was conducted to determine if the combination of traits presented in **Table 2** (excluding phenology) were able to explain a percentage of variation of BM_PM, HI, TGW, GM2, RUE_E40InB, RUE) InBA7, RUE_GF and RUET (**Table 3**). In total, 13.2% of the variation in final biomass (BM_PM) was explained by the combination of water index 2 (WI_2) and plant senescence reflectance index (PRSI). For HI, 18% of the variation was explained by the combination of LMA and chl*a* (RARSa) and 27.7% of the variation was explained when the model also considered chl*b*b (RARSb) and total chlorophyll content (R750_700.) In the case of TGW and GM2, the combination of enhanced vegetation index (EVI) and Plant Senescence Reflectance Index (PSRI) explained 13.3% and 16.5% of the variation, respectively, with opposite effects. RARSb explained 7.7% of the total variation observed in RUE_E40InB. However, the combination of LMA, Green Normalized Difference Vegetation Index (GNDVI), R750_700 and PSRI explained 25.6% of the variation in RUE_E40InB. For RUE_InBA7and RUE_GF, only 8.3% and 4% of the variation was explained by the combination of chlorophyll content in the flag leaf (SPAD_A7) and water index 4 (WI_4) or WI_2, respectively. Carotenoid content (RARSc) was the first component selected in the model and explained 3% of the variation in RUET. When structural independent pigment index (SIPI), WI_2 and NLamA7 were added to the model, 13.5% of RUET was explained.

**Table 3.** Stepwise analysis with biomass at physiological maturity (BM_PM), harvest index (HI), thousand grain weight (TGW), grain number (GM2), Radiation Use Efficiency measured between 40 days after emergence and initiation of booting (RUE_E40InB), RUE between initiation of booting and seven days after anthesis (RUE_InBA7), RUE between seven days after anthesis and physiological maturity (RUE_GF) and RUE between 40 days after emergence and physiological maturity (RUET) as dependent variables and nitrogen content and hyperspectral indices as independent variables.

| Trait | Variable chosen | *r* | *R*<sup>2</sup> | Significance |
| --- | --- | --- | --- | --- |
| BM_PM | WI_2 | 0.343a | 0.112 | <0.001 |
|  | WI_2, PSRI | 0.379b | 0.132 | <0.001 |
| HI | LMA(—) | 0.309a | 0.089 | <0.001 |
|  | LMA(—), RARSa | 0.437b | 0.180 | <0.001 |
|  | LMA(—), RARSa, RARSb | 0.502c | 0.236 | <0.001 |
|  | LMA(—), RARSa, RARSb, R750_700(—) | 0.544d | 0.277 | <0.001 |
|  | LMA(—), RARSa, RARSb, R750_700(—), RNDVI | 0.594e | 0.330 | <0.001 |
| TGW | EVI | 0.234a | 0.048 | <0.01 |
|  | EVI, PSRI | 0.380b | 0.133 | <0.001 |
| GM2 | EVI(—) | 0.304a | 0.086 | <0.001 |
|  | EVI(—), PSRI(—) | 0.420b | 0.165 | <0.001 |
| RUE_E40InB | RARSb | 0.289a | 0.077 | <0.001 |
|  | RARSb, LMA(—) | 0.345b | 0.107 | <0.001 |
|  | RARSb, LMA(—), GNDVI | 0.386c | 0.131 | <0.001 |
|  | LMA(—), GNDVI | 0.384d | 0.136 | <0.001 |
|  | LMA(—), GNDVI, R750_700(—) | 0.429e | 0.168 | <0.001 |
|  | LMA(—), GNDVI, R750_700(—), PSRI(—) | 0.525f | 0.256 | <0.001 |
| RUE_InBA7 | SPAD_A7(—) | 0.262a | 0.062 | 0.001 |
|  | SPAD_A7(—), WI_4 | 0.309b | 0.083 | 0.001 |
| RUE_GF | WI_2 | 0.214a | 0.039 | <0.01 |
| RUET | RARSc | 0.190a | 0.030 | <0.05 |
|  | RARSc, SIPI(—) | 0.270b | 0.060 | <0.01 |
|  | RARSc, SIPI(—), WI_2 | 0.341c | 0.098 | <0.001 |
|  | RARSc, SIPI(—), WI_2, NLamA7(—) | 0.397d | 0.135 | <0.001 |

### Genome-wide association

Marker trait association analyses carried out using best linear unbiased estimators (BLUEs) from two repetitions for each measured trait over two years. From across 23 traits 47 MTAs were identified with a -Log P value of 5 (P <0.00001) of which 10 passed FDR threshold determined in GAPIT (-Log P 7.12) for traits including total chlorophyll content (**Figure 5**), chlorophyll b content and carotenoid content (**Figure 1**). A full list of MTAs and plots can be found in **Table 4** and **SF9** respectively. Subgenome B had the most MTAs with 24 followed by the A and D genome with 11 found on each. The highest number of MTAs on a single chromosome were seen on 2B and 3B. The size of associated intervals varied greatly, ranging from less than 1Mbp to greater than 100Mbp with associations towards the centromere often being larger, consistent with the increase in centromeric linkage group size in wheat.

**Figure 5.**
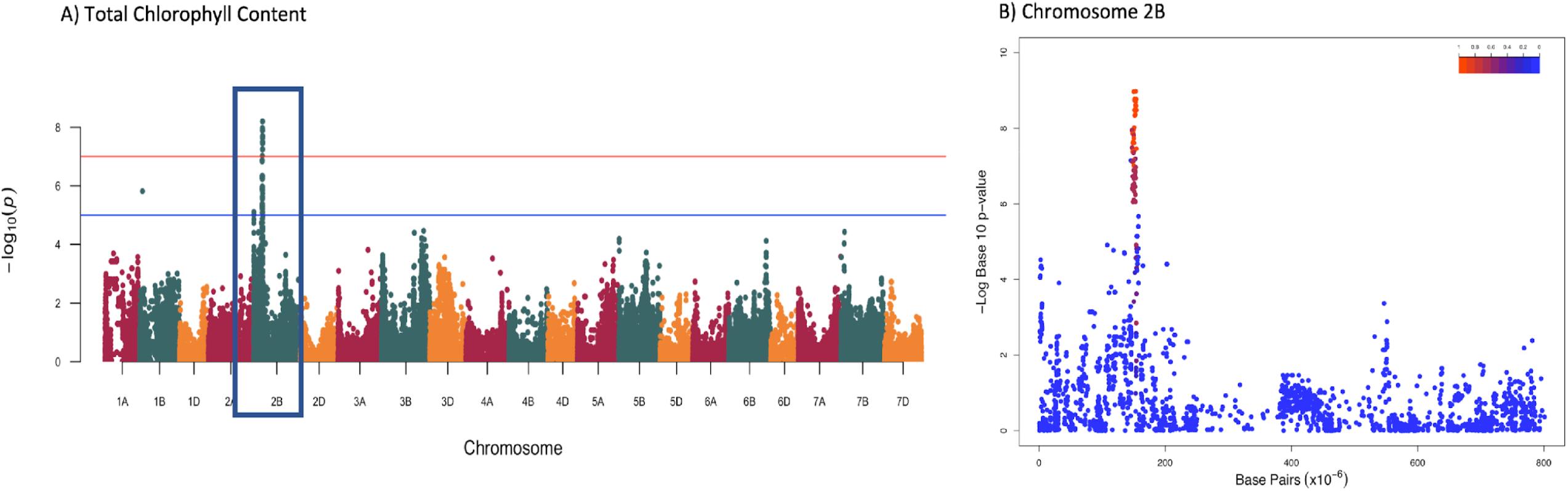
Genome-wide association results for total chlorophyll content. Manhattan plot showing A) the GWA output for total chlorophyll content (R750/700), significance cut-offs for -log10p of 5 and FDR correction are shown as blue and red lines respectively. B) The same GWA output for chromosome 2B, the level of genetic linkage to the most associated SNP is depicted by a heatmap.

**Table 4.** Summary of Marker-Trait Associations (MTAs) identified. Common SNPs are indicated by the same colour.

| Trait | Chromosome | MTA ID | Position | P-value | Interval |
| --- | --- | --- | --- | --- | --- |
| **Phenology** |  |  |  |  |  |
| DTA | chr5B | chr5B-27404243 | 27404243 | 8.12E-06 | 25-28Mbp |
|  | chr6B | chr6B-189683325 | 189683325 | 7.45E-07 | 188-189Mbp |
| **Nitrogen content** |  |  |  |  |  |
| SPAD | chr2B | chr2B-106930486 | 106930486 | 7.54E-06 | 105-140Mbp |
|  | chr2D | chr2D-16845994 | 16845994 | 1.96E-06 | 15.3-16.9Mbp |
|  | chr5A | chr5A-3550988 | 3550988 | 4.72E-07 | 3-4Mbp |
|  | chr6D | chr6D-456495062 | 456495062 | 8.69E-07 | 455-462Mbp |
| **Vegetation Index** |  |  |  |  |  |
| NDVI | chr7D | chr7D-608810464 | 608810464 | 2.56E-07 | 608.8-610Mbp |
| GNDVI | chr2B | chr2B-153275048 | 153275048 | 7.03E-10 | 148-157Mbp |
|  | chr3B | chr3B-723108033 | 723108033 | 2.11E-06 | 721-725Mbp |
| RNDVI | chr7A | chr7A-711982572 | 711982572 | 2.30E-06 | 711-712.5Mbp |
|  | chr7D | chr7D-608810464 | 608810464 | 9.92E-08 | 604-611Mbp |
| NDII | chr6A | chr6A-497983531 | 497983531 | 6.33E-06 | 497.8-498.5Mbp |
| NDMI | chr2B | chr2B-669669845 | 669669845 | 2.73E-07 | 530-670Mb |
| EVI | chr6B | chr6B-174740483 | 174740483 | 1.10E-06 | 17-19Mbp |
|  | chr3D | chr3D-523811772 | 523811772 | 1.50E-06 | 51-52Mbp |
| **Pigmentation composition** |  |  |  |  |  |
| RARSa | chr2A | chr2A-16347452 | 16347452 | 7.35E-08 | 500kb-26Mbp |
|  | chr2B | chr2B-20043565 | 20043565 | 8.67E-07 | 15-21Mbp |
|  | chr2B | chr2B-149018684 | 149018684 | 4.08E-08 | 148-154Mbp |
|  | chr7B | chr7B-135775607 | 135775607 | 5.78E-06 | 135-136Mbp |
| PSSa | chr2A | chr2A-17001351 | 17001351 | 7.13E-07 | 500kb-20Mbp |
| RARSb | chr2B | chr2B-153898371 | 153898371 | 1.83E-06 | 150-154Mbp |
|  | chr3B | chr3B-20186940 | 20186940 | 1.27E-07 | 18.4-21Mbp |
|  | chr3B | chr3B-715184359 | 715184359 | 9.06E-07 | 713-728Mbp |
| RARSc | chr3B | chr3B-20358957 | 20358957 | 1.25E-08 | 19-21.5Mbp |
|  | chr7A | chr7A-676592398 | 676592398 | 1.84E-06 | 675-677Mbp |
|  | chr7B | chr7B-722589303 | 722589303 | 4.66E-06 | 722-725Mbp |
|  | chr7D | chr7D-608810464 | 608810464 | 1.05E-06 | 608-609Mbp |
| SIPI | chr3B | chr3B-711119921 | 711119921 | 2.87E-06 | 690-712Mbp |
|  | chr7D | chr7D-610551080 | 610551080 | 3.74E-06 | 608-611Mbp |
| R750550 | chr2B | chr2B-2619111 | 2619111 | 9.55E-06 | 2.4-3.2Mbp |
|  | chr2B | chr2B-153275048 | 153275048 | 6.29E-09 | 150-155Mbp |
| R750700 | chr2B | chr2B-153275048 | 153275048 | 1.04E-09 | 150-155Mbp |
|  | chr3B | chr3B-736709296 | 736709296 | 2.72E-07 | 722-737Mbp |
| **Senescence/degradation indices** |  |  |  |  |  |
| NPQI | chr7A | chr7A-539405222 | 539405222 | 2.31E-07 | 523-564Mbp |
| PSRI | chr3B | chr3B-718670251 | 718670251 | 3.43E-07 | 715..7-718.6 |
| **Water Indices** |  |  |  |  |  |
| WI2 | chr3A | chr3A-616891767 | 616891767 | 9.04E-07 | 616-625Mbp |
| WI3 | chr1B | chr1B-572799923 | 572799923 | 1.09E-07 | 567-573Mbp |
|  | chr7A | chr7A-34727282 | 34727282 | 7.30E-06 | 34.5-35 Mbp |
| WI4 | chr3A | chr3A-616891767 | 616891767 | 2.87E-08 | 616-628Mbp |
|  | chr3D | chr3D-611549913 | 611549913 | 1.24E-07 | 612-620Mbp |
| **Physiology predictor** |  |  |  |  |  |
| LMA | chr3D | chr3D-336170196 | 336170196 | 3.29E-07 | 307-336Mbp |
|  | chr1B | chr1B-65889406 | 65889406 | 5.71E-07 | 65-83Mbp |
| R_DM | chr1D | chr1D-397474458 | 397474458 | 1.27E-06 | 396-399Mbp |

### Putative Candidate Genes and Haplotype Phased Non-Synonymous Variation

Candidate gene searches were carried out using Knetminer to identify genes within MTA intervals with phenotype/ontology terms associated with each trait alongside literature searches. Candidates were identified with ontology terms relating chlorophyll content and chlorophyll biosynthesis within multiple MTA intervals including Genome Uncoupled 5 (GUN5) in SPAD-2B, Early Chloroplast Biogenesis 2 (ECB1)/Vanilla Cream 1 (VAC1) in the shared interval RARSa-2b, RARSb-2b, R750/700-2B and R750/500-2B, SWEET4/5 bi-directional sugar transporter and Ethylene-responsive element binding factor 1 (ERF1) within 1Mbp of the MTA in RARSa-2A and PSSRa-2A. We also identified multiple candidates chlorophyll breakdown and senescence: protein phosphatase 2A (PP2A) and *HY5* a bZIP transcription factor that binds to the promoters of light-inducible genes in NPQI-7A, TCP20 and HK3 cytokinin receptor in PSRA-3B. Along with candidates that link to carotenoid biosynthesis and distribution: SYTF and TRAESCS3B02G039600 in RARSc-3B, TRAESCS7D02G503400 in RARSc-7D and also Small ubiquitin-related modifier 1 (SUMO1) in SIPI-3B and KNAT3/KNOTTED1-like in SIPI-7D, both of which have also been implicated in chlorophyll levels in the leaf.

The SWEET bidirectional sugar transporter was identified in the interval for chla content (RARSa) on chromosome 2A, whose closest orthologues are atSWEET4/5. These genes have been observed to have an effect on chlorophyll content in both *Arabidopsis thaliana^30^* and in *rice^31^*. A search was carried out for non-synonymous SNP calls within SNP calls that were removed prior to GWA, including many resulting from “off target” sequencing that did not have coverage in >90% panel members. This search identified a non-synonymous SNP at the start of the SWEET gene causing a substitution of Serine to Threonine. This variation was identified in 15 of 26 members with the minor allele of the MTA, suggesting this non-synonymous SNP is in phase with the MTA. No candidate genes that had any link to measured traits were identified for 4 MTAs. A full list of candidate genes can be seen in **ST6**.

## Discussion

### “De novo” SNP discovery

The size of the wheat genome means it is not yet economically viable to perform whole genome sequencing for large populations. Much of our understanding of the genetic diversity of wheat has come from array based genotyping^32,33^. An alternative is enrichment capture, that has uncovered “hidden variation” across world diversity panels^34^ and landraces^3^. Here we utilise capture sequencing and *de novo* SNP discovery to assess the contribution of strategically integrated exotic germplasm to overall CIMMYT germplasm diversity, to identify and track wild relative introgressions and demonstrate how this novel diversity can be exploited to identify candidate genes for agriculturally important traits.

### Recent synthetic wheat contribution to D genome variation and identification of wild relative introgressions

HiBAP panel members containing exotic pedigree history were found to have a 62% increase in genetic variation when compared to elite lines. Since this variation is present in wheat lines that have been demonstrated to have no yield penalty, it could be deployed rapidly into breeding programmes to alleviate the genetic bottleneck on the D genome which may be hindering genetic gains in wheat^35^. A large contribution to this variation is made by the synthetic derivatives in which the proportion of *Ae. tauschii* was observed to be as high as 43% of the D genome with an average of 16% across the synthetic derived subpopulation (**ST4**). Contribution of synthetic material to advanced lines has been theoretically predicted utilising genetic inference from pedigree history and allele frequency variation of markers at low resolution, determining a likely average of 17.5% contribution^8^. Here we are able to confirm these predictions and build on this by identifying regions to high resolution and through SNP identity utilising the *Ae. tauschii* reference genome. Across this whole subpopulation a total of 2.65Gb (67.2%) of the D genome was identified to be of synthetic donor origin, demonstrating the value of the HiBAP panel as a resource that could be utilised to study the effects of the presence of some of the ~22,000 genes found in these regions that are already introgressed into agronomically favourable backgrounds. We also identify that the proportion of *Ae. tauschii* donors in the synthetic subpopulation can vary substantially, from as little as 0.5% to as high as 43% of the D subgenome, a larger range than identified in studies using synthetic octoploids as primary *Ae. tauschii* donors (0.075 % to 13.5 %)^36^. Through comparison of sister lines within the panel we also show that the content of the donor genome is not necessarily linked to the number of subsequent crosses post introduction of primary synthetic material (**SF6, ST4**).

By utilising genome wide variation in conjunction with pedigree history and genetic resources for wheat wild relatives we are able to track introgressions from other common donors including Rye and *T. ponticum*. Using identification by genetic identity we tracked the once common CIMMYT 1BL/1RS introgression to 6 HiBAP panel members (**ST5**). We also confirm that this original introgression remains intact in each instance, demonstrating the inability of homologous recombination when the effect of ph mutants is negated, restricting breakage of alien introgressions^37^. We also identified a region of increased SNP density on chromosome 7DL in 3 panel members, pedigree history examination determined this to be of *T. ponticum* origin. This introgression, inferring resistance to both stem rust (Sr25) and leaf rust (Lr19)^17^ can be tracked through CIMMYT breeding material. The agronomic advantages these introgressions infer highlight the importance of tracking their presence in breeding programmes and for making selections for future crosses.

### Using high density genotyping and high-throughput phenotyping to uncover novel markers and putative candidate genes associated with photosynthetic efficiency

Recent developments in high-throughput phenotyping have already made significant contributions to physiological breeding^18,38,39^ and breeding programmes^40^. Assessment of photosynthetic related traits using high-throughput surrogates based on spectral profiles has allowed the identification of genetic variation in wheat for photosynthetic capacity and efficiency^24^ and respiration^20^ at leaf level along with pigmentation composition and water content at canopy level allowing the identification of QTLs associated with spectral indices^41,42^. In the present study, we aimed to identify genetic variation at leaf level from the spectral profiles reflected by green tissue. Our GWAS analysis identified 47 novel MTAs (**Table 4**) that, after a validation process, could be deployed into CIMMYT marker-assisted breeding programmes to track favourable alleles for traits that have important agronomic implications. One such trait is the chlorophyll content of leaves that has significant effect on photosynthetic efficiency, water use efficiency and yield in multiple crop species in field environments^43,44^. MTAs were identified for chl*a* and chl*b* separately and also for total chlorophyll content on chromosomes 2BS and 3BL with the main association on 2BS at ~150Mbp being present in all measurements which could be a result of the mostly overlapping absorbance spectra of the two pigments^45^. A QTL has been identified for chlorophyll content on 2BS under heat stress, spanning nearly the entire short arm of the chromosome^46^, using ultra-high density genotyping we are able to reduce this to an interval of <5Mbp. Within this interval we identify Early Chloroplast Biogenesis 2 (ECB1)/Vanilla Cream 1 (VAC1), a pentatricopeptide repeat protein with arabidopsis mutants in the gene reducing chlorophyll content by 10 fold^47,48^. SPAD measurements in the flag leaf (a surrogate of total chl content), uncovered MTAs on chromosomes 2B, 2D, 5A and 6D. Candidate gene search within the 2B interval showed the Genome Uncoupled 5 gene (GUN5). GUN5 encodes a magnesium chelatase ^49^, mutations in the gene in arabidopsis result in the disruption of chlorophyll synthesis. This can be explained because GUN 5 sits in the retrograde signalling pathway linking chlorophyll biosynthesis to sugar signalling^50^.

This makes GUN5 a possible target for future study in relation to chlorophyll variation in the HiBAP panel. We also identified a pigment specific MTA for chla on 2A and 7B and an MTA specific to chl*b* on chromosome 3BL. Of these, we believe that 3BL association has been identified previously in a QTL spanning over a fifth of 3B^51^, here we refine that to a region of less than 2Mbp. Within 1Mbp of MTA for chl*a* in 2A two candidate genes were identified: an ERF1 gene associated with biotic/abiotic stress tolerance, which when overexpressed in wheat has demonstrated a 50% increase in chl*a* content under normal growing conditions^52^. Also a SWEET bi-directional sugar transporter related to *Arabidopsis* SWEET4/5, with mutants in SWEET4/5affecting leaf chlorophyll content in *Arabidopsis*^30^ and rice^31^, although the mechanism for this effect is still unclear. SWEET transporters catalyse the passive efflux of sucrose down the gradient from the mesophyll to the apoplast where it is taken up by sucrose transporters (SUTs) into the phloem^53^. Within this gene, a non-synonymous mutation was identified in the prefiltered SNPs set that is in phase with the top SNP for this MTA, the SNP was called in 15 of the 25 lines containing the minor allele for this MTA. When taken together this evidence makes the SWEET gene a strong candidate for further study. The fact that both SWEET and GUN5 genes were associated with chlorophyll traits suggests that the two may be working together to control chlorophyll content in wheat leaves. Through the index NPQI we can also make inference about the level of chlorophyll degradation happening in the leaves^54^ which can be indicative of plant senescence. We detected an association for NPQI in chromosome 7AL that contains HY5, an antagonist of PIF that controls the accumulation of chlorophyll in the leaves in response to phytochrome photoreceptor signalling^55^, which also causes delayed senescence in rice transgenic plants^56^. We also observed an association on 3BL for plant senescence itself through calculation of PSRI, this interval contains AHK2 a cytokinin receptor which controls leaf longevity in Arabidopsis and knockouts demonstrate a delayed senescence phenotype. Another gene in this interval found >180Kbp from this MTA is transcription factor TCP20 which causes early senescence phenotypes in Arabidopsis but only in a double mutant including TCP9 due to predicted redundancy in signalling roles^57^.

Increasing RUE is a major target for achieving yield potential^21^. However, under field conditions like the ones experienced in Cd. Obregon, leaves are exposed to high irradiance absorbing more light they can use. This leads to photooxidative damage and reduces photosynthetic efficiency in a process known as photoinhibition. To avoid oxidative stress and photoinhibition, photoprotective mechanisms can be activated in response to high irradiance^58^. There is evidence that photoinhibition has a large impact on biomass production in crops exposed to high light levels^59^ and photoprotective mechanisms can increase yield and canopy RUE in rice^60^. Previous studies identified photoprotective genes associated with RUE^22^ indicating that protection of photosynthetic machinery has an impact in wheat. In this study, the photoprotective mechanisms that were detected are related with non-photochemical processes happening before photolysis in PSII, such as the xanthophyll cycle, where xanthophyll carotenoid plays an important role^59^. In our analysis, carotenoid content in the flag leaf (RARSc) was the first trait explaining 3% of RUET variation and together with carotenoids:chl*a* ratio (SIPI) 6% of variation was explained. Carotenoids play an important role protecting the photosynthetic machinery from excessive light^61,62^. Also, the ratio of carotenoids to chlorophyll is associated with senescence triggered by aging or stress^63^. An MTA for carotenoid content was identified on chromosome 3B, along with three others on homeologous regions of 7A, B and D. Two genes in the 3B interval were denoted as being involved in carotenoid biosynthesis by Knetminer, SYTF and TRAESCS3B02G039600.

### Reduced total chlorophyll content in the flag leaf is associated with enhanced RUE in elite cultivars

Total chlorophyll content was negatively correlated with RUE (**Table 3**) and it is mainly determined by chl*a* concentration since the ratio between chl*a*:chl*b* in wheat non-stressed plants is ~3:1^28^. In the present study, chl*a* (RARSa) explained a significant part of HI variation while chl*b* (RARSb) explained variation for RUE_E40InB. In wheat canopies, the uppermost layer including the flag leaf measured in this study receives the highest irradiance. However, photosynthetic machinery in wheat is saturated at 1,200 μmol m^-2^ s^-164^. Considering the high irradiance in Obregon when the measurements were taken (>1,800 μmol m^-2^ s^-1^), flag leaves absorb more light than they can use and need to engage photoprotective mechanisms. Lower chlorophyll content per unit area in the upper leaves facilitates light penetration in the canopy decreasing canopy extinction coefficient and therefore mitigating efficiency losses associated with light saturation^65,66^. This is in agreement with the negative effect observed here between total chl content and RUE, suggesting that less chlorophyll content in upper leaves has a positive effect on biomass production as observed for rice or soybean^43,44^.

### Implications in physiological breeding

Here we uncover the contribution of exotic material to variation in CIMMYT wheat lines, confirming the value of strategic incorporation of primary synthetics to ease genetic bottlenecks in the D genome. The HiBAP panel now represents an unprecedented resource of characterised genetic diversity, containing ~67% of the *Ae. tauschii* genome across the panel already in agronomically viable backgrounds. Identification of *Ae. tauschii* introgressions at such resolution allows breeders to select lines for strategic crossing to increase overall genetic variation using agronomically favourable material. The HiBAP panel has also been extensively phenotyped including yield traits, biomass accumulation, RUE and respiratory rates, in both yield potential and under abiotic stress. The results presented here highlight the possibilities created when phenotyping and genotyping efforts are coordinated in consortiums such as the International Wheat Yield Partnership (www.iwyp.org) to boost wheat genetic gains.

The use of hyperspectral reflectance (ASD Field Spec) measured on leaves in the field is independent of sunlight (due to the light source of the device) and one leaf can be measured in less than 30 seconds^19^. From the reflectance spectrum produced, multiple indices were derived which when combined with high density genotyping, facilitated the identification of candidate genes / traits integral to photosynthetic improvement. This protocol facilitates measurement of hundreds of genotypes per day to explore genetic variation of photosynthesis^24^ making association analysis feasible. The new MTAs for traits that contributed to RUE, HI and other traits of interest can be further used to identify allelic variation in other mapping populations or introgressed into elite lines through conventional and strategic crossing.

The current study was conducted on the flag leaf and scaling leaf-level data to the canopy can be complicated as leaf age and leaf angle can play a crucial role when integrating leaf level photosynthetic traits at the canopy scale^67^. However, leaf level measurements have been previously associated with higher yields^68–70^ and in the present study, traits measured at leaf level explained a significant proportion of the variation of BM_PM, HI, TGW, GM2 and RUE measured at different growth stages. Nevertheless, further experiments expanding measurements at the flag leaf to other canopy levels and at more time points across the whole growth cycle may be worthwhile^71^.

The identification of new sources of variation that contribute to increased photosynthetic potential in wheat together with the identification of markers associated with them could help to identify better donors that can provide superior combinations of alleles of useful genes.

## Materials and Methods

### Plant material

The High Biomass Association Mapping Panel (HiBAP) consists of 149 bread wheat spring types (**ST6**) that are agronomically acceptable including elite high yielding lines, pre-breeding lines that have been selected for high yield and/or biomass, including lines that have “exotic” material such as landraces or synthetic primary hexaploids in their recent pedigree history along with appropriate local check lines. The panel contains members that show broad variation of both RUE and biomass at multiple growth stages as described in^22^, and are controlled for the confounding effects of the extremes of height and phenology.

### DNA extraction and capture enrichment

Flag leaf tissue was obtained from plants used in field trials after anthesis. Material from 10 individuals was taken per line and pooled for DNA extraction using a standard CTAB based method. DNA purity was assessed using a NanoDrop 2000 (Thermofisher Scientific) and quantified fluorometrically using the Quant-iTTM assay kit (Life Technologies). Dual indexed DNA libraries were constructed with a modal insert size of 450bp using TruSeq DNA library preparation kit (Illumina). Capture enrichment was carried out using the MyBaits targeted capture kit (Arbor Bioscience, Michigan USA) incorporating 100,000 custom 120-mer RNA bait sequences with 8x precapture multiplexing following standard protocols. In total, 90,000 probes were designed in an island strategy distributed to facilitate enrichment and subsequent variant calling from regions spanning the entire genome. Probe sequences were designed based on a subgenome-collapsed reference, allowing probes to target homeologous regions, increasing the genomic design space with the fewest probes possible. 10,000 probes were designed to target selected gene sequences using an end-to-end tiling strategy covering the gene body and the promoter region (~2000bp). The 2 Kb distance was based on the median distance between the TSS and the first transposable element, 1.52 Kb to allow a high likelihood of full promoter sequence capture^72^. Enriched libraries were then sequenced on a NovaSeq6000 (Illumina) S4 flowcell producing 2 x 150bp paired end sequences.

### Genotyping and imputation

Sequencing quality was assessed with FastQC and low-quality reads removed/trimmed. The paired-end sequencing data for each accession was mapped to the Refseq-v1.0 reference sequence^77^ using BWA MEM version 0.7.13^73^. Mapping results were filtered using SAMtools v1.4^74^; any non-uniquely mapping reads, unmapped reads, and/or poor-quality reads were removed. PCR duplicates were identified and removed using Picard Tools MarkDuplicates. Variant calling was carried out using bcftools and were filtered using GATK^75^, using the standard GATK recommended parameters of minimum quality of 30, a minimum depth of 5. The likely functional effect of each variant was annotated using SnpEff 4.3^76^ using a custom database generated using Refseq v1.1 annotation^77^. For each SNP loci found in the panel as a whole, if no alternative allele was found for an individual but mapping depth of ≥5 was observed, the individual was designated as homozygous reference for that loci, else it was designated as missing data. SNP loci that had <10% missing data and a minor allele frequency of ≥5% were then subjected to imputation using Beagle 5.0 ^78^

### Population structure analysis

Genetic inference into the population structure of the panel was made using STRUCTURE 2.3.4^79^ using model based Bayesian approaches along with Hierarchical clustering to deduce similarity between lines. An admixture model was selected in STRUCTURE and run using 30,000 burn-in iterations and 50,000 repetitions of the Markov Chain Monte Carlo (MCMC) model for the assumed sub-populations of k (2-10) for 10 independent, randomly seeded iterations of the analysis per assumed sub-population. To identify the statistically most likely number of definable subpopulations, the delta k method of^80^ was applied to all 10 replicates, where the Dk statistic is deduced from the rate of change in the probability of likelihood [LnP(D)] value between each k value. The Evanno method was implemented through STRUCTURE HARVESTER Python script^81^. CLUMPP 1.1.2 was used to produce a consensus Q matrix using 10 independent STRUCTURE replicates for each assumed subpopulation number^82^. To assess the overall level of genetic variation in the panel PCA analysis was carried out using the Scikit-Learn machine learning package in Python. PCA was applied to genotyping data from the 35K wheat breeders array and capture enrichment genotyping data for all subgenomes combined and for the D genome separately to assess the level of variation that could be observed when using pre-known SNP loci and using *de novo* methods. To determine genomic regions most greatly affected by the incorporation of exotic material, fixation index (Fst) was calculated in windows of 500Kbp in pairwise comparisons of the elite subpopulation with the exotic subpopulations that included landrace, synthetic and landrace+synthetic and the introgression lines subpopulation as a whole. As a control measure, the elite population was randomly split and Fst was calculated between the pseudo subpopulations.

### Identification of Ae. tauschii synthetic D genome donor regions and S. cereale introgressions

To identify genomic regions originating from *Ae. tauschii* donors used in the creation of primary synthetics present in the pedigree history of 40 HiBAP panel members, SNPs called for each HiBAP member were compared to SNPs called between the CS wheat reference and the *Ae. tauschii* reference genome^83^ to determine identity. Paired end 150bp reads were simulated from the *Ae. tauschii* reference genome to a depth of 20x using WGSim. Reads were mapped and variant called using the same methods outlined for the capture sequencing of the HiBAP panel members. To remove noise created by varietal SNPs between CIMMYT germplasm and the CS reference, Weebill1 SNPs were generated using trimmed sequencing reads used to create the contig assembly of Weeblil1 (project PRJEB35709 accession SAMEA6374024) and SNPs from the panel matching in location and allele we removed from further analysis. SNPs across each panel member were binned into 500Kbp bins, within each bin the number of SNPs matching *Ae. Tauschii* variation was counted. To estimate the contribution of *Ae. tauschii* within the entirety of the HiBAP panel the maximum number of SNPs within every 500kbp window was assessed for both the elite background and synthetic background subpopulations. Where the value for a bin in any synthetic line was 5-fold higher than the average value for each bin from the whole elite population this was classed as a modern *Ae. tauschii* region. Additionally, the theoretical contribution to pedigree from *Ae. tauschii* was estimated using the available pedigree and selection history information from the International Wheat Information System (IWIS), curated by CIMMYT.

To determine the presence of Rye introgressions in the panel SNPs between Rye and CS were generated by mapping and variant calling Lo7 Rye Illumina sequencing reads (ERS446995) against CS as previously described. Genetic identification of regions of Rye was carried out using the same methods for *Ae. Tauschii* outlined above.

### Field experimental conditions

The 149 lines were grown in two consecutive growing seasons (2015/16 and 2016/17, which will be referred to as Y16 and Y17 respectively). Field experiments were carried out at the IWYP-Hub (International Wheat Yield Partnership Phenotyping Platform) at CENEB in the Yaqui Valley, near Ciudad Obregón, Sonora, México (27°24’ N, 109°56’ W, 38 masl), under fully irrigated conditions.

The soil type was a coarse sandy clay, mixed montmorillonitic typic caliciorthid, low in organic matter, and slightly alkaline (pH 7.7) in nature^84^. Experimental design was a α-lattice with four replications in raised beds (2 beds per plot each 0.8 m wide) with four (Y16) and two (Y17) rows per bed (0.1 m and 0.24 m between rows respectively) and 4 m long. The emergence dates were 7 Dec. 2015 and 30 Nov. 2016 for each year. The seeding rate was 102 Kg ha^-1^. Appropriate fertilization, weed disease and pest control were implemented to avoid yield limitations. Plots were fertilized with 50 kg N ha^-1^ (urea) and 50 kg P ha^-1^ at soil preparation, 50 kg N ha^-1^ with the first irrigation and another 150 kg N ha^-1^ with the second irrigation. Growing conditions and main agronomic characteristics of the trial grown for two years are detailed in^22^.

### Field traits measurements

Agronomical and physiological characterization was conducted as detailed in^22^. In summary, two replicates were measured for the different traits with the exception of phenology, yield, thousand grain weight (TGW) and grain number per m^2^ (GNO) where four replicates were scored.

Anthesis (DTA) and physiological maturity (DTM) dates were recorded using the scale for growth stages (GS) developed by^85^, following the average phenology of the plot when 50% of the shoots reached GS65 and GS87, respectively. The phenological stages represent the number of days from emergence to the onset of these stages.

At physiological maturity, a sub-sample consisting of 100 (Y16) or 50 (Y17) fertile shoots was obtained from the harvested area to estimate yield components and harvest index (HI). Grain yield was determined in 3.2 – 4 m^2^ using standard protocols^86^ discarding 50cm in both extremes of the plots to avoid edge effects arising from extra solar radiation reaching border plants. A subsample of grains was weighed before and after drying (oven dried at 70°C for 48 h) and the ratio of dry to fresh weight was used to determine dry grain yield and for TGW.

In order to calculate radiation use efficiency (RUE) at different growth stages, different biomass samplings were collected forty (Y16) or forty-two (Y17) days after emergence (BME40), at initiation of booting (BMInB), approximately seven days after anthesis (BMA7) and after physiological maturity (BM_PM). BME40, BMInB and BMA7 consisted of all aboveground tissue from two central beds of a 50-cm length, starting at least 50 cm from the end of the plot (or the previous harvest) to avoid border effects. Fresh biomass was oven-dried at 70°C for 48 h for dry weight measurement. BM_PM was calculated from yield/HI. RUE was then estimated as the slope of linear regression of cumulative aboveground biomass on cumulative intercepted PAR^87^ considering the different biomass sampling such as RUE_E40InB: from 40 days after emergence to initiation of booting, RUE_InBA7: from initiation of booting to seven days after anthesis, RUE_GF: RUE from seven days after anthesis until physiological maturity and RUET: RUE from 40 days after emergence to physiological maturity. Light intercepted by the canopy was not used for the RUE calculations due to low heritability estimates and the lack of genotypic effect^22^. The genotypes evaluated in this study intercept an average of 80% at E40, 95% at InB and 94% at A+7^22^. We assumed that 100% of the radiation was intercepted between E40 and A+7 and 50% was intercepted for one quarter of the grain filling period^88^.

### Chlorophyll content, N composition and reflectance measurements

Chlorophyll content in the flag leaf was measured with a SPAD-502 Minolta (Spectrum Technologies Inc., Plainfield, IL, USA) in five flag leaves per plot seven days after anthesis (SPADA7).

Nitrogen concentration of the leaf lamina (NlamA7) was measured using the Kjeldahl digestion method putting together all green leaf laminas from 12 random stems harvested seven days after anthesis after drying (oven dried at 70°C for 48 h), milling and digestion with concentrated sulphuric acid.

The hyperspectral reflectance of flag leaves was measured between 11.00 to 14.00 hours approximately seven days after anthesis following the protocol described by^19^. A FieldSpec^®^3 (Analytical Spectral Devices, Boulder, CO, USA) full range spectroradiometer (350–2500 nm) was coupled via a fibre optic cable to a leaf. A mask was used to reduce the leaf-clip aperture and a black circular gasket was pasted to the mask to avoid leaf damage and to eliminate potential entry of external light through the edges. One reflectance measurement was made per leaf lamina, and two measurements per plot in two plots per entry. Different spectral reflectance indices were calculated as described by^89–96^. The formulas for index calculations are presented in **Table 2**. Leaf mass area (LMA) and respiration on a dry matter basis (RDM) were estimated using a web-application to predict wheat physiological traits from hyperspectral reflectance spectra known as the *Wheat Physiology Predictor* [https://www.metabolome-express.org/pheno/] based on^19^ and^20^ prediction models.

### Genome-wide association analysis

Association analysis was carried out using GAPIT^97^ on 149 HiBAP lines. A model based on the unified mixed linear model approach, the SUPER algorithm, was applied to the genotype/phenotype data. The model was adjusted using membership coefficient matrices produced by STRUCTURE assuming between 2-8 subpopulations (Q2-8) or the first 10 eigenvectors from principal component analysis (PC1-10) along with a kinship matrix (K) as covariates to limit the confounding effects of population structure effects and therefore reducing false positives. The EMMA method proposed by^98^ to create a positive semidefinite kinship matrix (K) was followed, implemented in GAPIT. Interval size was determined by taking the flanking SNPs from each association that were greater than the lower t -Log P value of 5 threshold. To identify possible causative candidates, genes within the associated intervals were submitted to Knetminer^99^. The resultant information networks were assessed and if adequate evidence was available to suggest the gene or its orthologous genes may be involved in a mechanism linking to the trait to which it was associated, the gene was selected as a possible candidate. Interval genes were also mined for non-synonymous variants in both the high confidence SNP calls along with those falling below depth filters.

### Statistical analysis of phenotypic data

Data for field measured traits were analysed by using a mixed model for computing the least square means (LSMEANS) for each genotype across both years using the program Multi Environment Trial Analysis with R for Windows (METAR,^100^). DTA was used as covariate (fixed effect) when its effect was significant with the exception of phenology and RUE. Broad sense heritability (*H*^2^) was estimated for each trait across both years as:

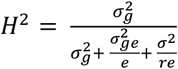

where r = number of repetitions, e= number of environments (years), σ^2^ =error variance, σ^2^g =genotypic variance and σ^2^ge = G×Y variance.

Multiple linear regression analysis (stepwise) was used to analyse the relationship between the studied variables using the SPSS statistical package (SPSS Inc., Chicago, IL, USA)

## Data availability

The datasets generated during during this study are available in the European Nucleotide Archive repository (www.ebi.ac.uk/ena/) under study name PRJEB38874.

## Acknowledgements

RJ, BC, LG and AH were supported by funding from the BBSRC and IWYP. RJ, LG and AH were supported by funding from the BBSRC and IWYP (BB/N020871/1; BB/P016855/1) and Designing Future Wheat. GM, FJPC, CRA and MR were supported by the Sustainable Modernization of Traditional Agriculture (MasAgro) initiative from the Secretariat of Agriculture, Livestock, Rural Development, Fisheries and Food (SAGARPA) and the International Wheat Yield Partnership (IWYP) project. We acknowledge the GWP Physiology team in Obregon for supporting field experiments management and data collection and Jacob Enk at Arbor Bioscience for assistance with capture development.

## Author contributions

AH, MR, JRE, RTF and GM conceptualised the project. RJ, BC and LJG carried out genotyping and genetic analyses. GM, CRA and FJPC carried out phenotypic measurements. RJ and GM wrote the manuscript. All authors edited and approved the manuscript.

## Bibliography

1. Haas, M., Schreiber, M. & Mascher, M. Domestication and crop evolution of wheat and barley: Genes, genomics, and future directions. J. Integr. Plant Biol. 61, 204–225 (2019).

2. Dvorak, J., Luo, M. C., Yang, Z. L. & Zhang, H. B. The structure of the Aegilops tauschii genepool and the evolution of hexaploid wheat. Theor. Appl. Genet. 97, 657–670 (1998).

3. Gardiner, L.-J. et al. Hidden variation in polyploid wheat drives local adaptation. Genome Res. 28, 1319–1332 (2018).

4. Rimbert, H. et al. High throughput SNP discovery and genotyping in hexaploid wheat. PLoS ONE 13, e0186329 (2018).

5. Das, M. K., Bai, G., Mujeeb-Kazi, A. & Rajaram, S. Genetic diversity among synthetic hexaploid wheat accessions (Triticum aestivum) with resistance to several fungal diseases. Genet. Resour. Crop Evol. 63, 1285–1296 (2016).

6. Jafarzadeh, J. et al. Breeding value of primary synthetic wheat genotypes for grain yield. PLoS ONE 11, e0162860 (2016).

7. Li, A., Liu, D., Yang, W., Kishii, M. & Mao, L. Synthetic hexaploid wheat: yesterday, today, and tomorrow. ENG. 4, 552–558 (2018).

8. Rosyara, U. et al. Genetic contribution of synthetic hexaploid wheat to cimmyt’s spring bread wheat breeding germplasm. Sci. Rep. 9, 12355 (2019).

9. Manès, Y. et al. Genetic Yield Gains of the CIMMYT International Semi-Arid Wheat Yield Trials from 1994 to 2010. Crop Sci. 52, 1543–1552 (2012).

10. Lopes, M. S. & Reynolds, M. P. Drought Adaptive Traits and Wide Adaptation in Elite Lines Derived from Resynthesized Hexaploid Wheat. Crop Sci. 51, 1617–1626 (2011).

11. Cossani, C. M. & Reynolds, M. P. Heat Stress Adaptation in Elite Lines Derived from Synthetic Hexaploid Wheat. Crop Sci. 55, 2719–2735 (2015).

12. Colmer, T. D., Flowers, T. J. & Munns, R. Use of wild relatives to improve salt tolerance in wheat. J. Exp. Bot. 57, 1059–1078 (2006).

13. Velu, G. et al. Genetic dissection of grain zinc concentration in spring wheat for mainstreaming biofortification in CIMMYT wheat breeding. Sci. Rep. 8, 13526 (2018).

14. Imtiaz, M., Ogbonnaya, F. C., Oman, J. & van Ginkel, M. Characterization of quantitative trait loci controlling genetic variation for preharvest sprouting in synthetic backcross-derived wheat lines. Genetics 178, 1725–1736 (2008).

15. Kishii, M. An update of recent use of aegilops species in wheat breeding. Front. Plant Sci. 10, 585 (2019).

16. Ren, T.-H. et al. Development and characterization of a new 1BL. 1RS translocation line with resistance to stripe rust and powdery mildew of wheat. Euphytica 169, 207–213 (2009).

17. Niu, Z. et al. Development and characterization of wheat lines carrying stem rust resistance gene Sr43 derived from Thinopyrum ponticum. Theor. Appl. Genet. 127, 969–980 (2014).

18. Araus, J. L. & Cairns, J. E. Field high-throughput phenotyping: the new crop breeding frontier. Trends Plant Sci. 19, 52–61 (2014).

19. Silva-Perez, V. et al. Hyperspectral reflectance as a tool to measure biochemical and physiological traits in wheat. J. Exp. Bot. 69, 483–496 (2018).

20. Coast, O. et al. Predicting dark respiration rates of wheat leaves from hyperspectral reflectance. Plant Cell Environ. 42, 2133–2150 (2019).

21. Zhu, X.-G., Long, S. P. & Ort, D. R. Improving photosynthetic efficiency for greater yield. Annu. Rev. Plant Biol. 61, 235–261 (2010).

22. Molero, G. et al. Elucidating the genetic basis of biomass accumulation and radiation use efficiency in spring wheat and its role in yield potential. Plant Biotechnol. J. 17, 1276–1288 (2019).

23. Driever, S. M., Lawson, T., Andralojc, P. J., Raines, C. A. & Parry, M. A. J. Natural variation in photosynthetic capacity, growth, and yield in 64 field-grown wheat genotypes. J. Exp. Bot. 65, 4959–4973 (2014).

24. Silva-Pérez, V. et al. Genetic variation for photosynthetic capacity and efficiency in spring wheat. J. Exp. Bot. 71, 2299–2311 (2020).

25. Blackburn, G. A. Hyperspectral remote sensing of plant pigments. J. Exp. Bot. 58, 855–867 (2007).

26. Filella, I., Serrano, L., Serra, J. & Peñuelas, J. Evaluating Wheat Nitrogen Status with Canopy Reflectance Indices and Discriminant Analysis. Crop Sci. 35, 1400–1405 (1995).

27. Evans, J. R. & Clarke, V. C. The nitrogen cost of photosynthesis. J. Exp. Bot. 70, 7–15 (2019).

28. Watanabe, N., Evans, J. R. & Chow, W. S. Changes in the photosynthetic properties of australian wheat cultivars over the last century. Functional Plant Biol. 21, 169 (1994).

29. Gardiner, L.-J. et al. Analysis of the recombination landscape of hexaploid bread wheat reveals genes controlling recombination and gene conversion frequency. Genome Biol. 20, 69 (2019).

30. Liu, X., Zhang, Y., Yang, C., Tian, Z. & Li, J. AtSWEET4, a hexose facilitator, mediates sugar transport to axial sinks and affects plant development. Sci. Rep. 6, 24563 (2016).

31. Zhou, Y. et al. Overexpression of OsSWEET5 in rice causes growth retardation and precocious senescence. PLoS ONE 9, e94210 (2014).

32. Wang, S. et al. Characterization of polyploid wheat genomic diversity using a high-density 90,000 single nucleotide polymorphism array. Plant Biotechnol. J. 12, 787–796 (2014).

33. Allen, A. M. et al. Characterization of a Wheat Breeders’ Array suitable for high-throughput SNP genotyping of global accessions of hexaploid bread wheat (Triticum aestivum). Plant Biotechnol. J. 15, 390–401 (2017).

34. Pont, C. et al. Tracing the ancestry of modern bread wheats. Nat. Genet. 51, 905–911 (2019).

35. Lopes, M. S. et al. Exploiting genetic diversity from landraces in wheat breeding for adaptation to climate change. J. Exp. Bot. 66, 3477–3486 (2015).

36. Nyine, M. et al. Understanding the introgression process from *Aegilops tauschii* into hexaploid wheat through identity by descent analysis and its effect on genetic diversity. BioRxiv (2019). doi:10.1101/855106

37. Hao, M. et al. The resurgence of introgression breeding, as exemplified in wheat improvement. Front. Plant Sci. 11, 252 (2020).

38. Chapman, S. et al. Pheno-Copter: A Low-Altitude, Autonomous Remote-Sensing Robotic Helicopter for High-Throughput Field-Based Phenotyping. Agronomy 4, 279–301 (2014).

39. Tattaris, M., Reynolds, M. P. & Chapman, S. C. A Direct Comparison of Remote Sensing Approaches for High-Throughput Phenotyping in Plant Breeding. Front. Plant Sci. 7, 1131 (2016).

40. Reynolds, M. et al. Breeder friendly phenotyping. Plant Science 110396 (2020).

41. Gizaw, S. A., Godoy, J. G. V., Pumphrey, M. O. & Carter, A. H. Spectral Reflectance for Indirect Selection and Genome-Wide Association Analyses of Grain Yield and Drought Tolerance in North American Spring Wheat. Crop Sci. 58, 2289 (2018).

42. Liu, C., Pinto, F., Cossani, C. M., Sukumaran, S. & Reynolds, M. Spectral reflectance indices as proxies for yield potential and heat stress tolerance in spring wheat: heritability estimates and marker-trait associations. (2019).

43. Slattery, R. A., VanLoocke, A., Bernacchi, C. J., Zhu, X.-G. & Ort, D. R. Photosynthesis, Light Use Efficiency, and Yield of Reduced-Chlorophyll Soybean Mutants in Field Conditions. Front. Plant Sci. 8, 549 (2017).

44. Gu, J. et al. Rice ( Oryza sativa L.) with reduced chlorophyll content exhibit higher photosynthetic rate and efficiency, improved canopy light distribution, and greater yields than normally pigmented plants. Field Crops Res. 200, 58–70 (2017).

45. Huang, J. et al. Meta-Analysis of the Detection of Plant Pigment Concentrations Using Hyperspectral Remotely Sensed Data. PLoS ONE 10, e0137029 (2015).

46. Bhusal, N., Sharma, P., Sareen, S. & Sarial, A. K. Mapping QTLs for chlorophyll content and chlorophyll fluorescence in wheat under heat stress. Biol. Plant. 62, 1–11 (2018).

47. Yu, Q.-B., Jiang, Y., Chong, K. & Yang, Z.-N. AtECB2, a pentatricopeptide repeat protein, is required for chloroplast transcript accD RNA editing and early chloroplast biogenesis in Arabidopsis thaliana. Plant J. 59, 1011–1023 (2009).

48. He, P. et al. GhYGL1d, a pentatricopeptide repeat protein, is required for chloroplast development in cotton. BMC Plant Biol. 19, 350 (2019).

49. Davison, P. A. & Hunter, C. N. Abolition of magnesium chelatase activity by the gun5 mutation and reversal by Gun4. FEBS Lett. 585, 183–186 (2011).

50. McCormac, A. C. & Terry, M. J. The nuclear genes Lhcb and HEMA1 are differentially sensitive to plastid signals and suggest distinct roles for the GUN1 and GUN5 plastid-signalling pathways during de-etiolation. Plant J. 40, 672–685 (2004).

51. Czyczyło-Mysza, I. et al. Quantitative trait loci for leaf chlorophyll fluorescence parameters, chlorophyll and carotenoid contents in relation to biomass and yield in bread wheat and their chromosome deletion bin assignments. Mol. Breed. 32, 189–210 (2013).

52. Xing, L. et al. Overexpression of ERF1-V from Haynaldia villosa Can Enhance the Resistance of Wheat to Powdery Mildew and Increase the Tolerance to Salt and Drought Stresses. Front. Plant Sci. 8, 1948 (2017).

53. Chen, L.-Q. et al. Sucrose efflux mediated by SWEET proteins as a key step for phloem transport. Science 335, 207–211 (2012).

54. Lowe, A., Harrison, N. & French, A. P. Hyperspectral image analysis techniques for the detection and classification of the early onset of plant disease and stress. Plant Methods 13, 80 (2017).

55. Toledo-Ortiz, G. et al. The HY5-PIF regulatory module coordinates light and temperature control of photosynthetic gene transcription. PLoS Genet. 10, e1004416 (2014).

56. Burman, N., Bhatnagar, A. & Khurana, J. P. OsbZIP48, a HY5 Transcription Factor Ortholog, Exerts Pleiotropic Effects in Light-Regulated Development. Plant Physiol. 176, 1262–1285 (2018).

57. Danisman, S. et al. Arabidopsis class I and class II TCP transcription factors regulate jasmonic acid metabolism and leaf development antagonistically. Plant Physiol. 159, 1511–1523 (2012).

58. Murchie, E. H. & Niyogi, K. K. Manipulation of photoprotection to improve plant photosynthesis. Plant Physiol. 155, 86–92 (2011).

59. Murchie, E. H., Ali, A. & Herman, T. Photoprotection as a trait for rice yield improvement: status and prospects. Rice (N Y) 8, 31 (2015).

60. Hubbart, S. et al. Enhanced thylakoid photoprotection can increase yield and canopy radiation use efficiency in rice. Commun. Biol. 1, 22 (2018).

61. Demmig-Adams, B. Carotenoids and photoprotection in plants: A role for the xanthophyll zeaxanthin. Biochimica et Biophysica Acta (BBA) - Bioenergetics 1020, 1–24 (1990).

62. Cazzonelli, C. I. Goldacre Review: Carotenoids in nature: insights from plants and beyond. Functional Plant Biol. 38, 833 (2011).

63. Bort J., Araus J.L. & Casadesús J. in Irrigation systems performance (eds. Bogliotti C., Lamaddalena N., Lebdi F. & Todorovic M.) 52, 251–263 (Bari: CIHEAM, 2005).

64. Blum, A. Variation among wheat cultivars in the response of leaf gas exchange to light. J. Agric. Sci. 115, 305–311 (1990).

65. Ort, D. R. & Melis, A. Optimizing antenna size to maximize photosynthetic efficiency. Plant Physiol. 155, 79–85 (2011).

66. Hamblin, J., Stefanova, K. & Angessa, T. T. Variation in chlorophyll content per unit leaf area in spring wheat and implications for selection in segregating material. PLoS ONE 9, e92529 (2014).

67. Furbank, R. T., Jimenez-Berni, J. A., George-Jaeggli, B., Potgieter, A. B. & Deery, D. M. Field crop phenomics: enabling breeding for radiation use efficiency and biomass in cereal crops. New Phytol. 223, 1714–1727 (2019).

68. Fischer, R. A. et al. Wheat Yield Progress Associated with Higher Stomatal Conductance and Photosynthetic Rate, and Cooler Canopies. Crop Sci. 38, 1467 (1998).

69. Carmo-Silva, E. et al. Phenotyping of field-grown wheat in the UK highlights contribution of light response of photosynthesis and flag leaf longevity to grain yield. J. Exp. Bot. 68, 3473–3486 (2017).

70. Tang, Y. et al. Yield, growth, canopy traits and photosynthesis in high-yielding, synthetic hexaploid-derived wheats cultivars compared with non-synthetic wheats. Crop Pasture Sci. 68, 115 (2017).

71. Murchie, E. H. et al. Measuring the dynamic photosynthome. Ann. Bot. 122, 207–220 (2018).

72. Wicker, T. et al. Impact of transposable elements on genome structure and evolution in bread wheat. Genome Biol. 19, 103 (2018).

73. Li, H. & Durbin, R. Fast and accurate short read alignment with Burrows-Wheeler transform. Bioinformatics 25, 1754–1760 (2009).

74. Li, H. et al. The Sequence Alignment/Map format and SAMtools. Bioinformatics 25, 2078–2079 (2009).

75. McKenna, A. et al. The Genome Analysis Toolkit: a MapReduce framework for analyzing next-generation DNA sequencing data. Genome Res. 20, 1297–1303 (2010).

76. Cingolani, P. et al. A program for annotating and predicting the effects of single nucleotide polymorphisms, SnpEff: SNPs in the genome of *Drosophila melanogaster* strain w1118; iso-2; iso-3. Fly (Austin) 6, 80–92 (2012).

77. International Wheat Genome Sequencing Consortium (IWGSC) et al. Shifting the limits in wheat research and breeding using a fully annotated reference genome. Science 361, (2018).

78. Browning, B. L. & Browning, S. R. Genotype Imputation with Millions of Reference Samples. Am. J. Hum. Genet. 98, 116–126 (2016).

79. Pritchard, J. K., Stephens, M. & Donnelly, P. Inference of population structure using multilocus genotype data. Genetics 155, 945–959 (2000).

80. Evanno, G., Regnaut, S. & Goudet, J. Detecting the number of clusters of individuals using the software STRUCTURE: a simulation study. Mol. Ecol. 14, 2611–2620 (2005).

81. Earl, D. A. & vonHoldt, B. M. STRUCTURE HARVESTER: a website and program for visualizing STRUCTURE output and implementing the Evanno method. Conserv. Genet. Resour. 4, 359–361 (2012).

82. Jakobsson, M. & Rosenberg, N. A. CLUMPP: a cluster matching and permutation program for dealing with label switching and multimodality in analysis of population structure. Bioinformatics 23, 1801–1806 (2007).

83. Luo, M.-C. et al. Genome sequence of the progenitor of the wheat D genome Aegilops tauschii. Nature 551, 498–502 (2017).

84. Sayre, K. D., Rajaram, S. & Fischer, R. A. Yield potential progress in short bread wheats in northwest mexico. Crop Sci. 37, 36–42 (1997).

85. Zadoks, J. C., Chang, T. T. & Konzak, C. F. A decimal code for the growth stages of cereals. Weed Res. 14, 415–421 (1974).

86. Pietragalla, J. Physiological Breeding II: A Field Guide to Wheat Phenotyping. (2012).

87. Monteith, J. L. & Moss, C. J. Climate and the Efficiency of Crop Production in Britain [and Discussion]. Philosophical Transactions of the Royal Society of London. Series B, Biological Sciences 281, 277–294 (1977).

88. Reynolds, M. P., van Ginkel, M. & Ribaut, J. M. Avenues for genetic modification of radiation use efficiency in wheat. J. Exp. Bot. 51 Spec No, 459–473 (2000).

89. Peñuelas, J., Filella, I., Biel, C., Serrano, L. & Savé, R. The reflectance at the 950-970 nm region as an indicator of plant water status. Int. J. Remote Sens. 14, 1887–1905 (1993).

90. Peñuelas, J. & Filella, I. Visible and near-infrared reflectance techniques for diagnosing plant physiological status. Trends Plant Sci. 3, 151–156 (1998).

91. Araus, J. L., Casadesus, J. & Bort, J. Recent tools for the screening of physiological traits determining yield. (2001).

92. Babar, M. A. et al. Spectral Reflectance to Estimate Genetic Variation for In-Season Biomass, Leaf Chlorophyll, and Canopy Temperature in Wheat. Crop Sci. 46, 1046 (2006).

93. Gutierrez, M., Reynolds, M. P. & Klatt, A. R. Association of water spectral indices with plant and soil water relations in contrasting wheat genotypes. J. Exp. Bot. 61, 3291–3303 (2010).

94. Lobos, G. A. et al. Wheat genotypic variability in grain yield and carbon isotope discrimination under Mediterranean conditions assessed by spectral reflectance. J. Integr. Plant Biol. 56, 470–479 (2014).

95. Penuelas, J., Baret, F. & Filella, I. Semi-Empirical Indices to Assess Carotenoids/Chlorophyll-a Ratio from Leaf Spectral Reflectance. Photosynthetica (1995).

96. Chappelle, E. W., Kim, M. S. & McMurtrey, J. E. Ratio analysis of reflectance spectra (RARS): An algorithm for the remote estimation of the concentrations of chlorophyll A, chlorophyll B, and carotenoids in soybean leaves. Remote Sensing of Environment 39, 239–247 (1992).

97. Lipka, A. E. et al. GAPIT: genome association and prediction integrated tool. Bioinformatics 28, 2397–2399 (2012).

98. Kang, H. M. et al. Variance component model to account for sample structure in genome-wide association studies. Nat. Genet. 42, 348–354 (2010).

99. Hassani-Pak, K. et al. Developing integrated crop knowledge networks to advance candidate gene discovery. Appl. Transl. Genomics 11, 18–26 (2016).

100. META-R (Multi Environment Trial Analysis with R for Windows) Version 6.04 - CIMMYT Research Software. at <https://data.cimmyt.org/dataset.xhtml?persistentId=hdl:11529/10201>

