## Supplementary Figures for "Uncovering candidate genes involved in photosynthetic capacity using unexplored genetic variation in Spring Wheat"

The following Supporting Information is available for this article:

**Table ST1.** DNA sequencing and mapping statistics
**Table ST2.** The distribution of SNPs called against the Refseq1.0 Chinese Spring wheat reference genome.
**Table ST3.** Genome wide distribution of SNPs used for GWA analysis
**Table ST4.** Identification of regions of *Ae. Tauschii* origin in HiBAP lines with Synthetic pedigree history
**Table ST5.** Lines in the HiBAP panel containing *S. cereale*/*T. ponticum* introgressions
**Table ST6.** Pedigree history information for the HiBAP panel
**Figure SF1**. Depiction of genotyping capture probe set design/tiling strategy
**Figure SF2.** A heatmap ideogram demonstrating genome wide SNP density across the HiBAP panel
**Figure SF3.** Estimation of the most likely number of true subpopulations (K_number) within the HiBAP panel
**Figure SF4.** Principal component analysis showing genetic variation in the HiBAP panel
**Figure SF5.** Genome wide Fst calculations between subpopulations within the HiBAP panel
**Figure SF6.** SNP density plot of two synthetically derived sister lines
**Figure SF7**. SNP density plots for chromosome 1B from each member of the HiBAP panel found to contain the 1BS/1RL Rye introgression
**Figure SF8**. SNP density plots for chromosome 7D from each member of the HiBAP panel found to contain the 7DL/7EL *Thinopyrum ponticum* introgression
**Figure SF9.** Manhattan plots showing GWA results for 23 traits.

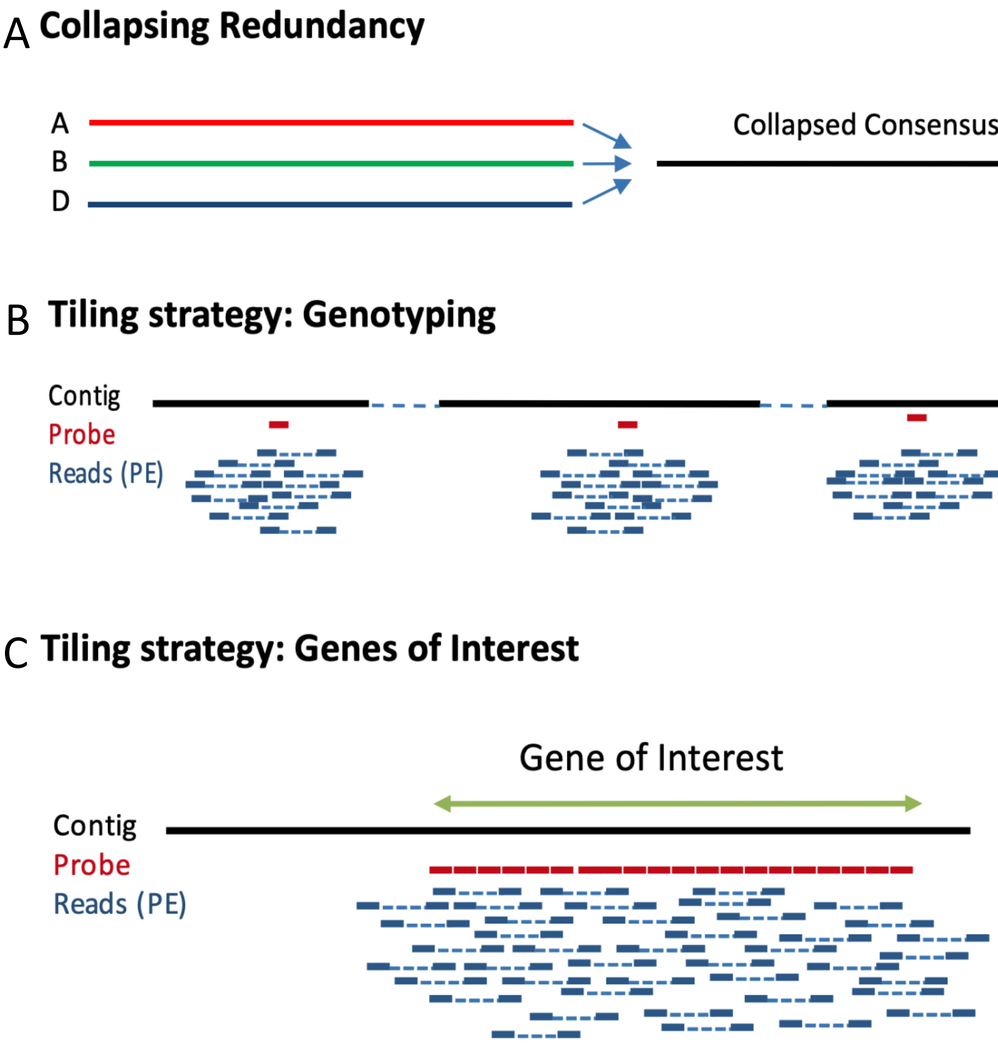


SF1: Depiction of genotyping capture probe set design/tiling strategy including A) collapsing the reference genome B) the "genotyping" portion as probes scattered across the genome C) with the "gene of interest" portion in which probes were arranged almost end to end across the gene body and promoter region.

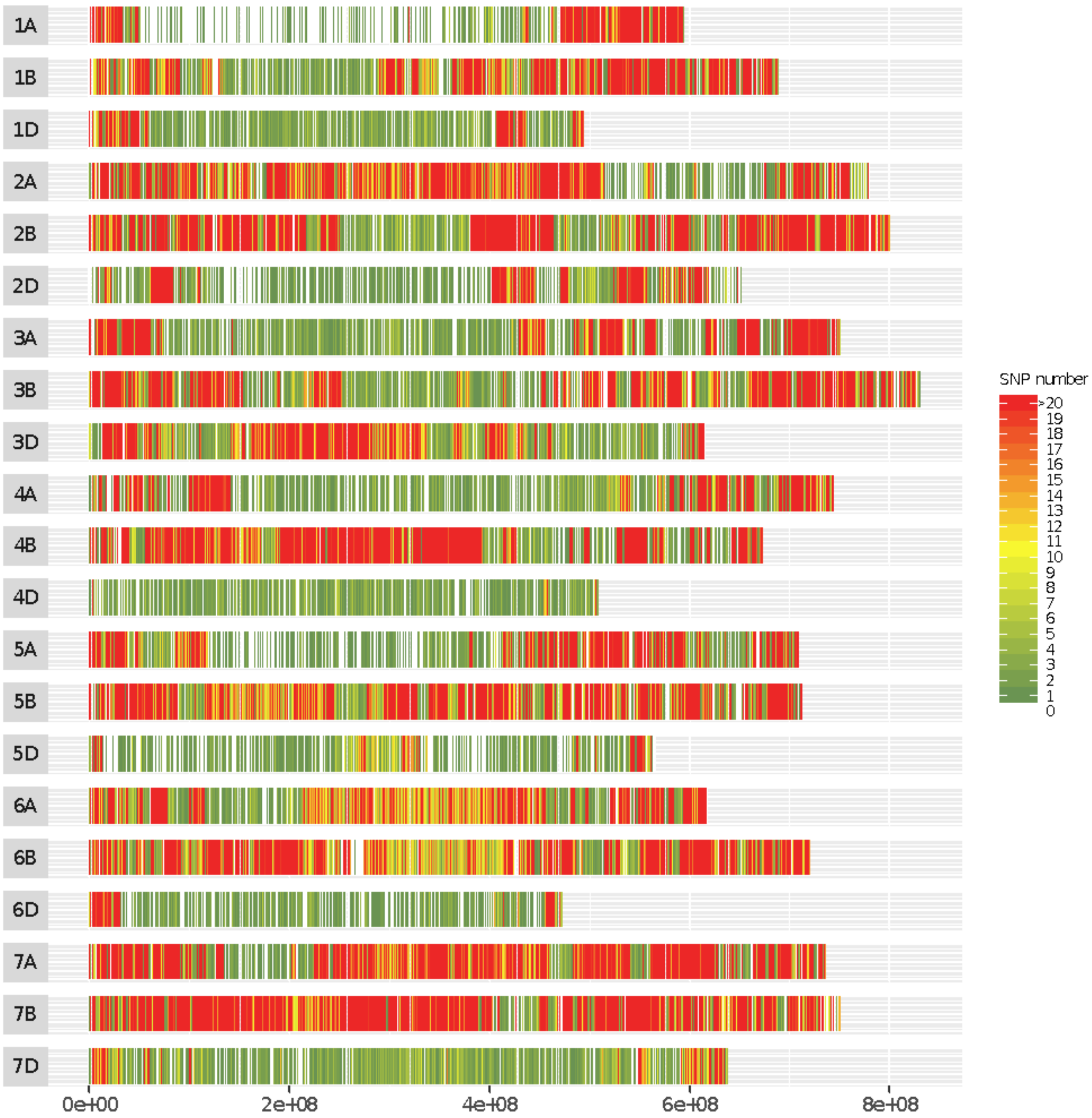


SF2: A heatmap ideogram demonstrating genome wide SNP density across the HiBAP panel after all called SNPs were combined and subsequently filtered to remove loci with >10% missing data and a minor allele frequency of less than 5%.

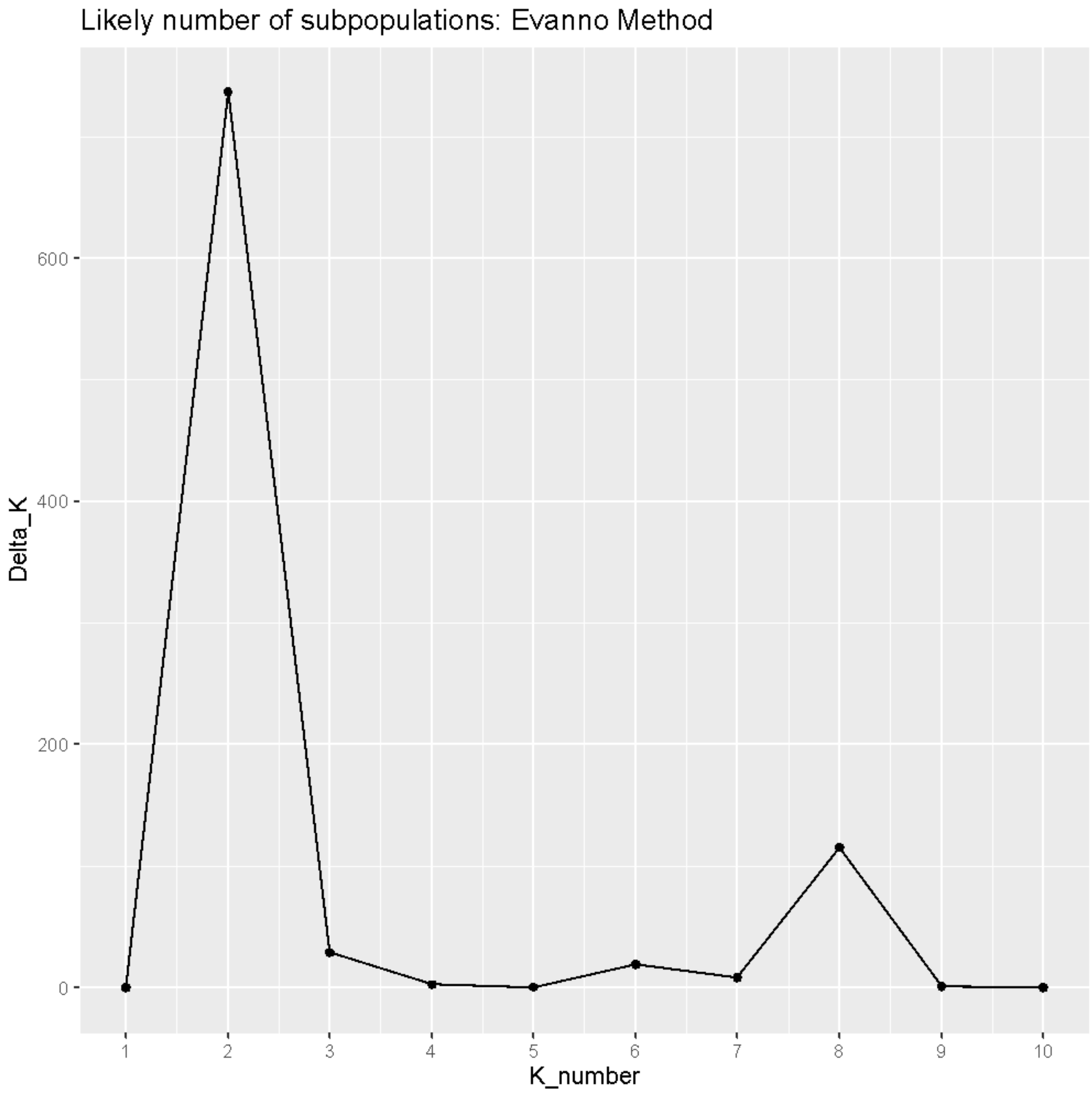

SF3: Estimation of the most likely number of true subpopulations (K_number) within the HiBAP panel estimated using the Evanno Method demonstrating the presence of 2 main subpopulations.

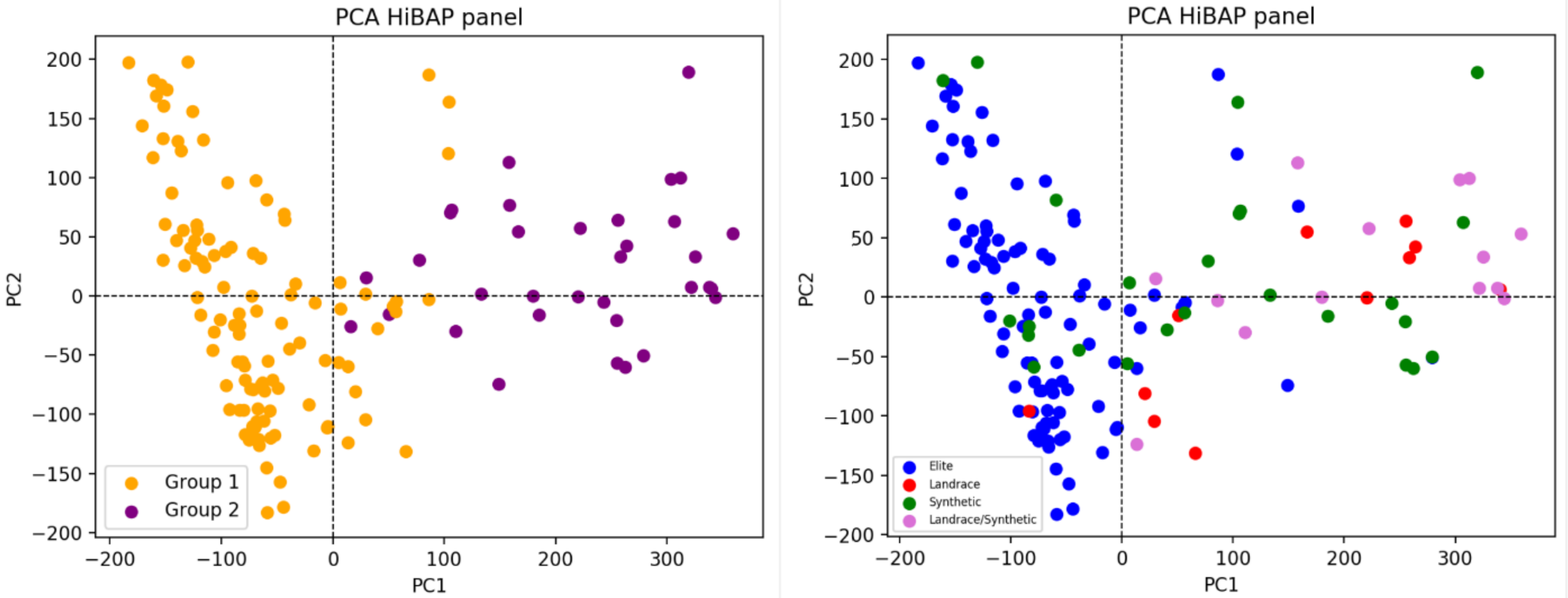


SF4: Principal component analysis showing genetic variation in the HiBAP panel. A) Panel members when coloured by subpopulation membership deduced using STRUCTURE software. B) Panel members coloured by presence of landrace, synthetic or a combination in their pedigree history.

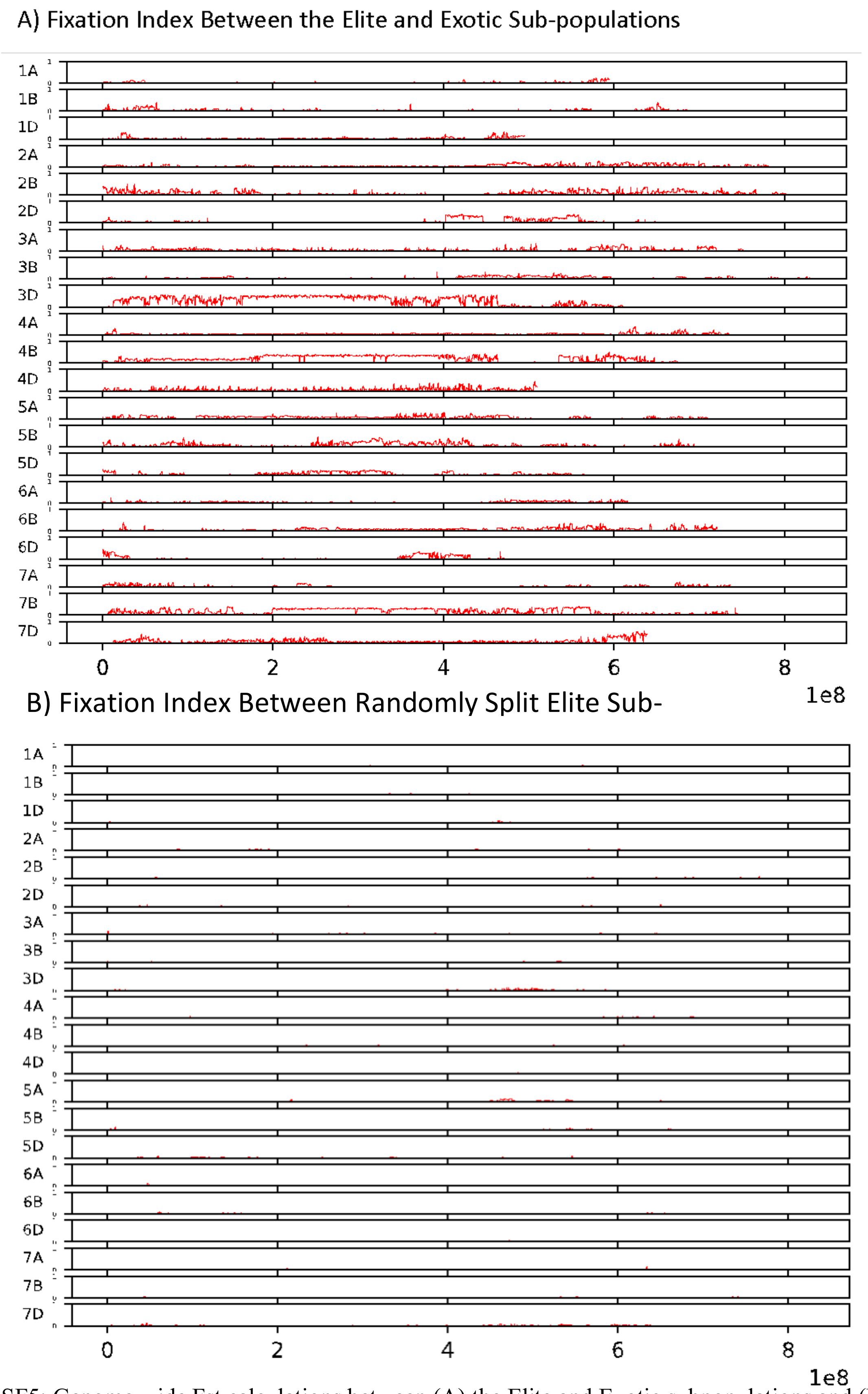


SF5: Genome wide Fst calculations between (A) the Elite and Exotic subpopulations and (B) two pseudo-subpopulations of elite background wheat created by randomly splitting the elite subpopulation into two groups.

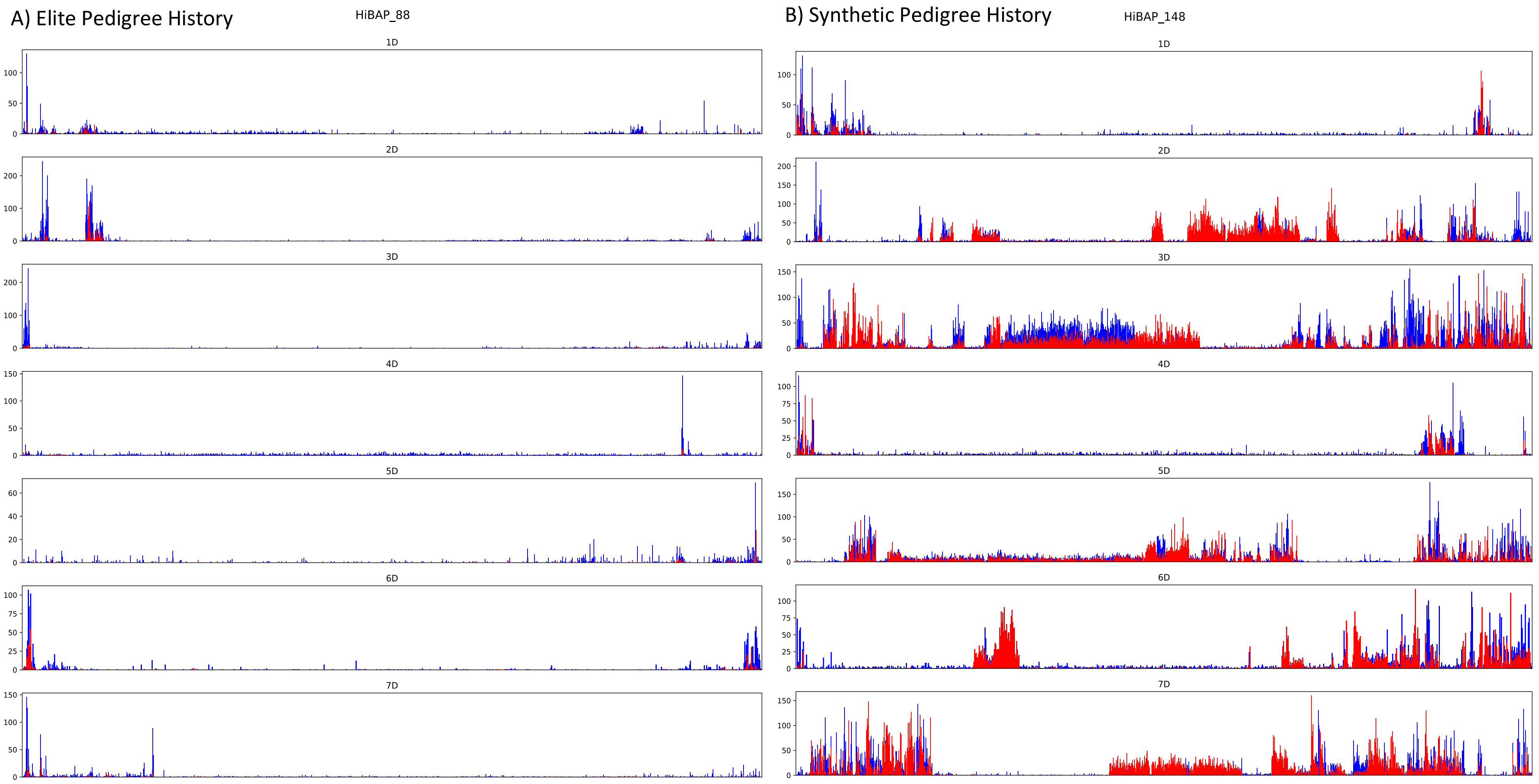


SF6: SNP density plot of two synthetically derived sister lines where SNPs are in 500Kbp bins. Red bars indicate number of SNPs identified to match modern Ae. Tauschii, blue bars show all other SNPs. Demonstrating the altering levels of introgressed regions, even between members of the same cross and selection history.

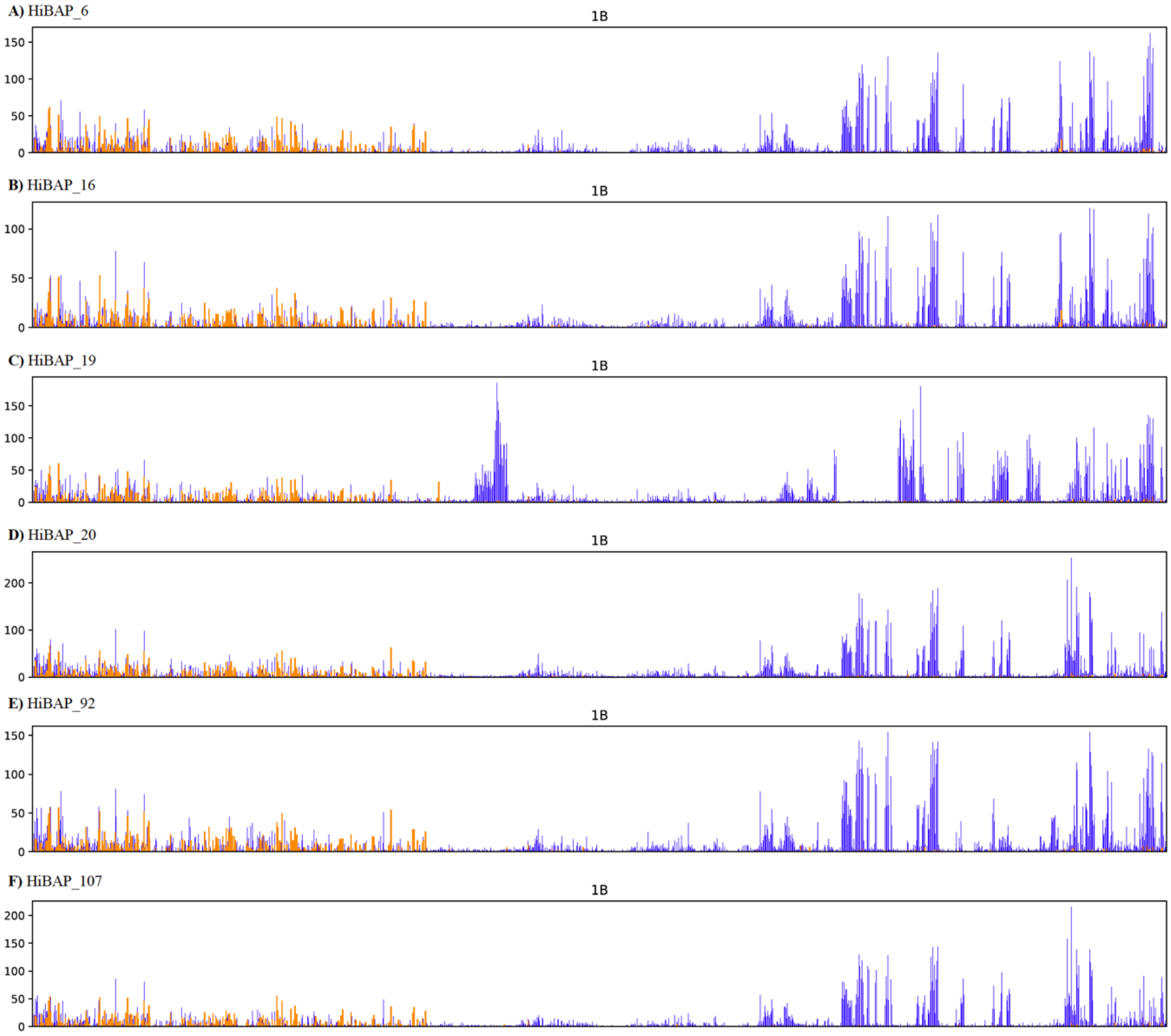


SF7: SNP density plots for chromosome 1B from each member of the HiBAP panel found to contain the 1BS/1RL Rye introgression. Orange bars indicate the number of SNPs per 500Kbp bin that match the position and allele with those found in between Rye and the wheat reference genome. Blue bars show number of SNPs per bin that do not match Rye genetic variation.

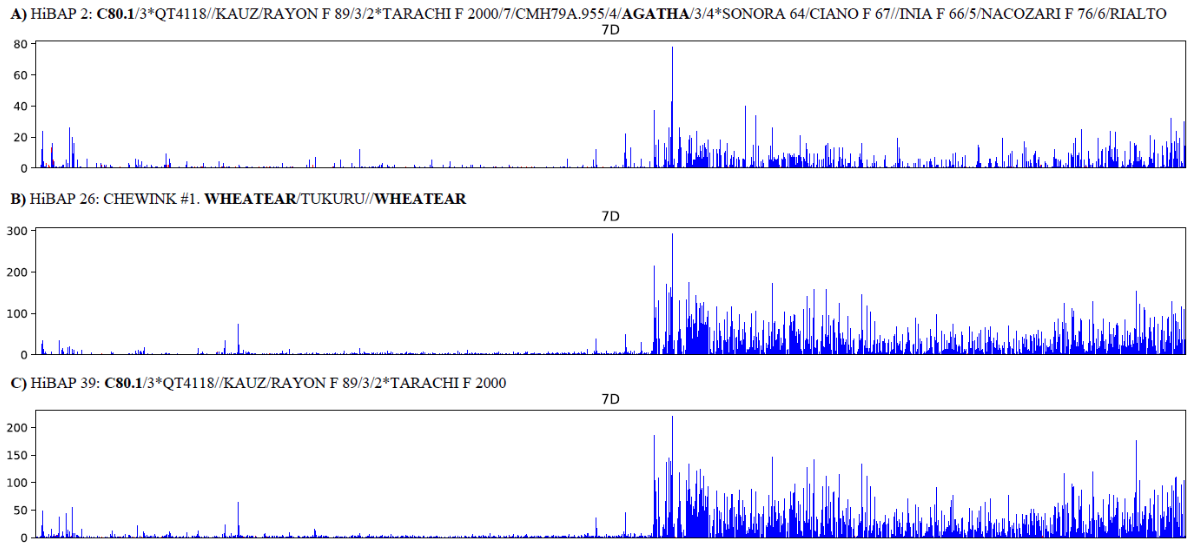


SF8: SNP density plots for chromosome 7D from each member of the HiBAP panel found to contain the 7DL/7EL *Thinopyrum ponticum* introgression. Blue bars show number of SNPs per 500Kbp bin.

SF9: Manhattan plots showing GWA results for 23 traits. Blue line depicts the significance threshold of -Log10 P 5 and the red line depicts the FDR threshold.

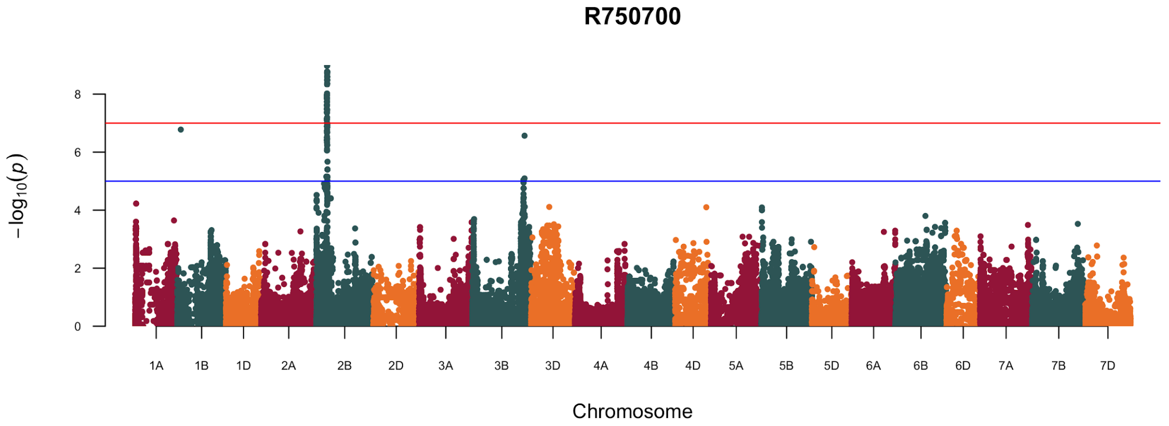


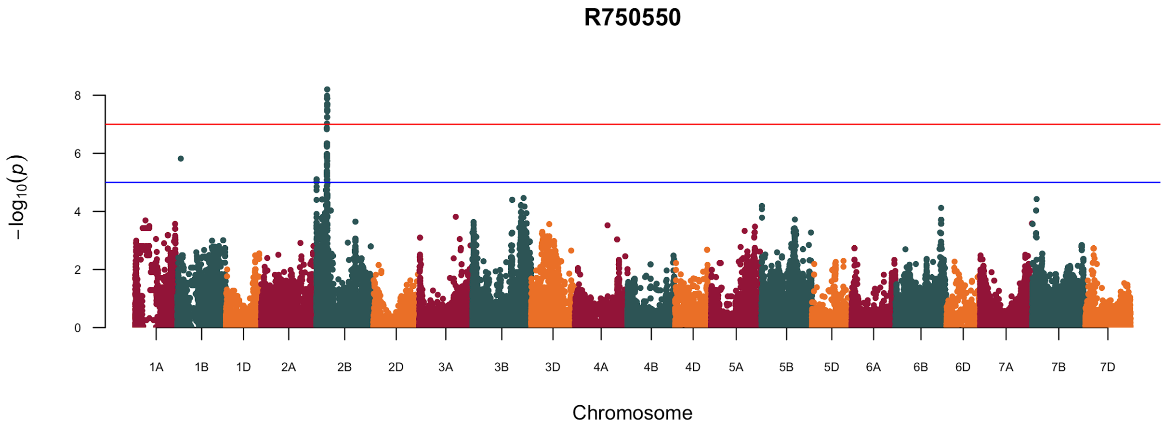


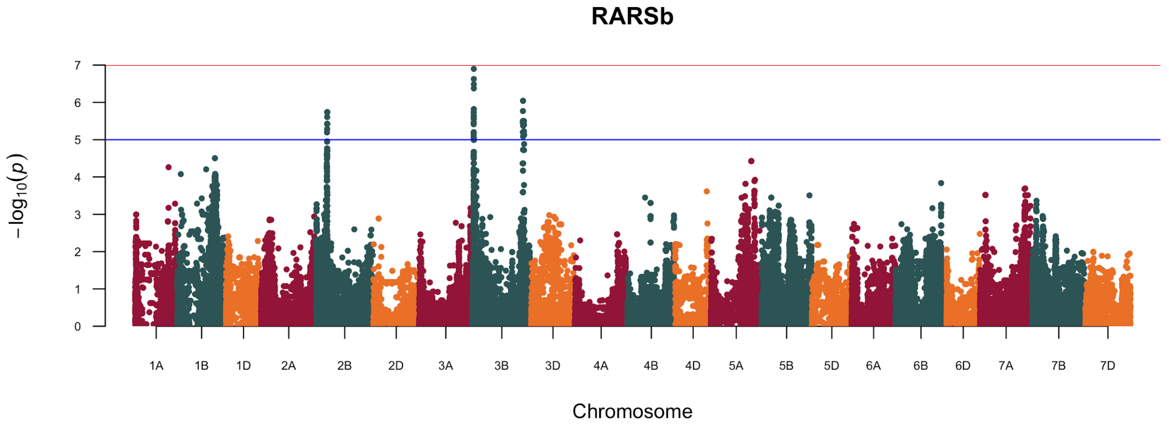

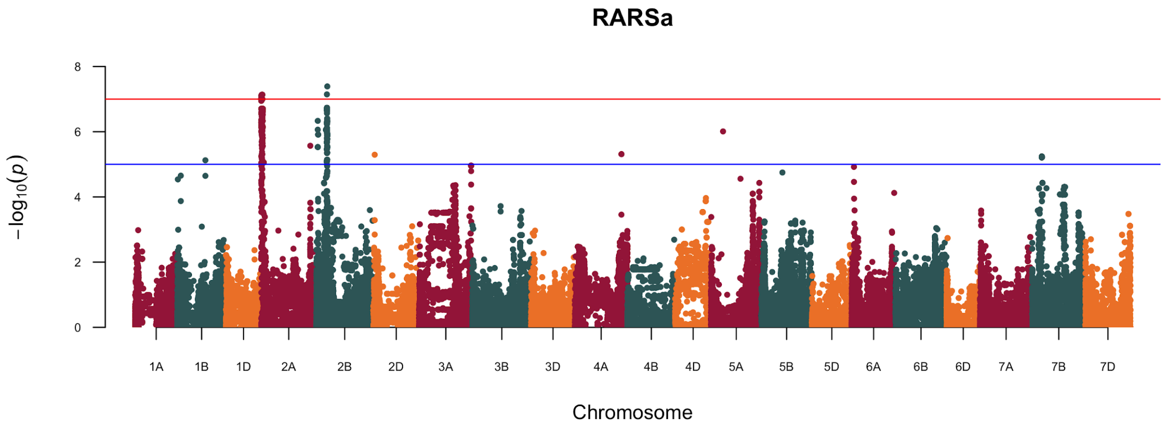
RARSa
−log10(p)
Chromosome
1A 1B 1D 2A 2B 2D 3A 3B 3D 4A 4B 4D 5A 5B 5D 6A 6B 6D 7A 7B 7D

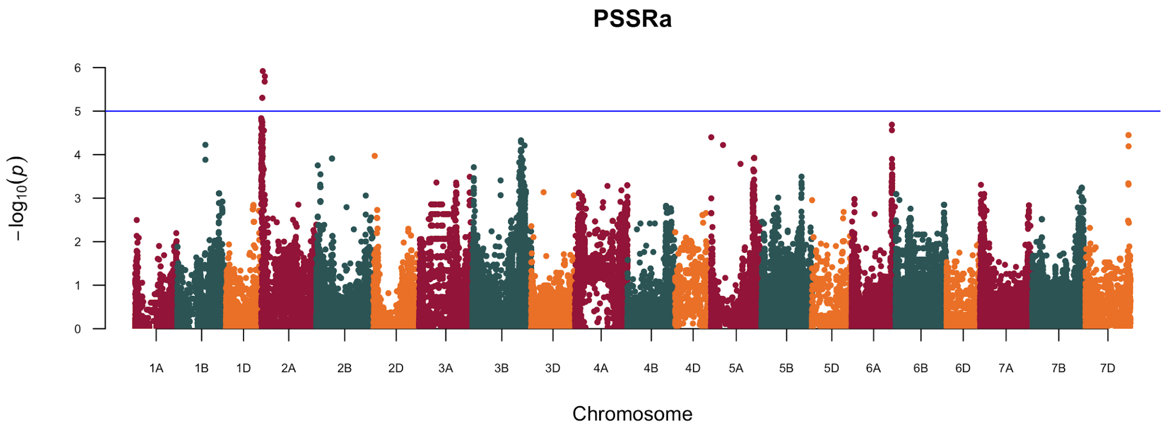
PSSRa
−log10(p)
Chromosome
1A 1B 1D 2A 2B 2D 3A 3B 3D 4A 4B 4D 5A 5B 5D 6A 6B 6D 7A 7B 7D

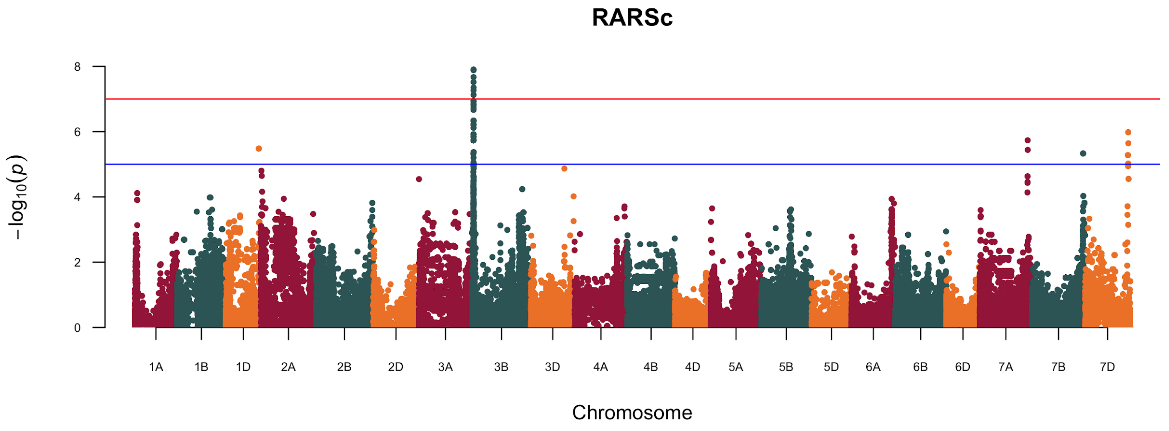
RARSc
−log10(p)
Chromosome
1A 1B 1D 2A 2B 2D 3A 3B 3D 4A 4B 4D 5A 5B 5D 6A 6B 6D 7A 7B 7D

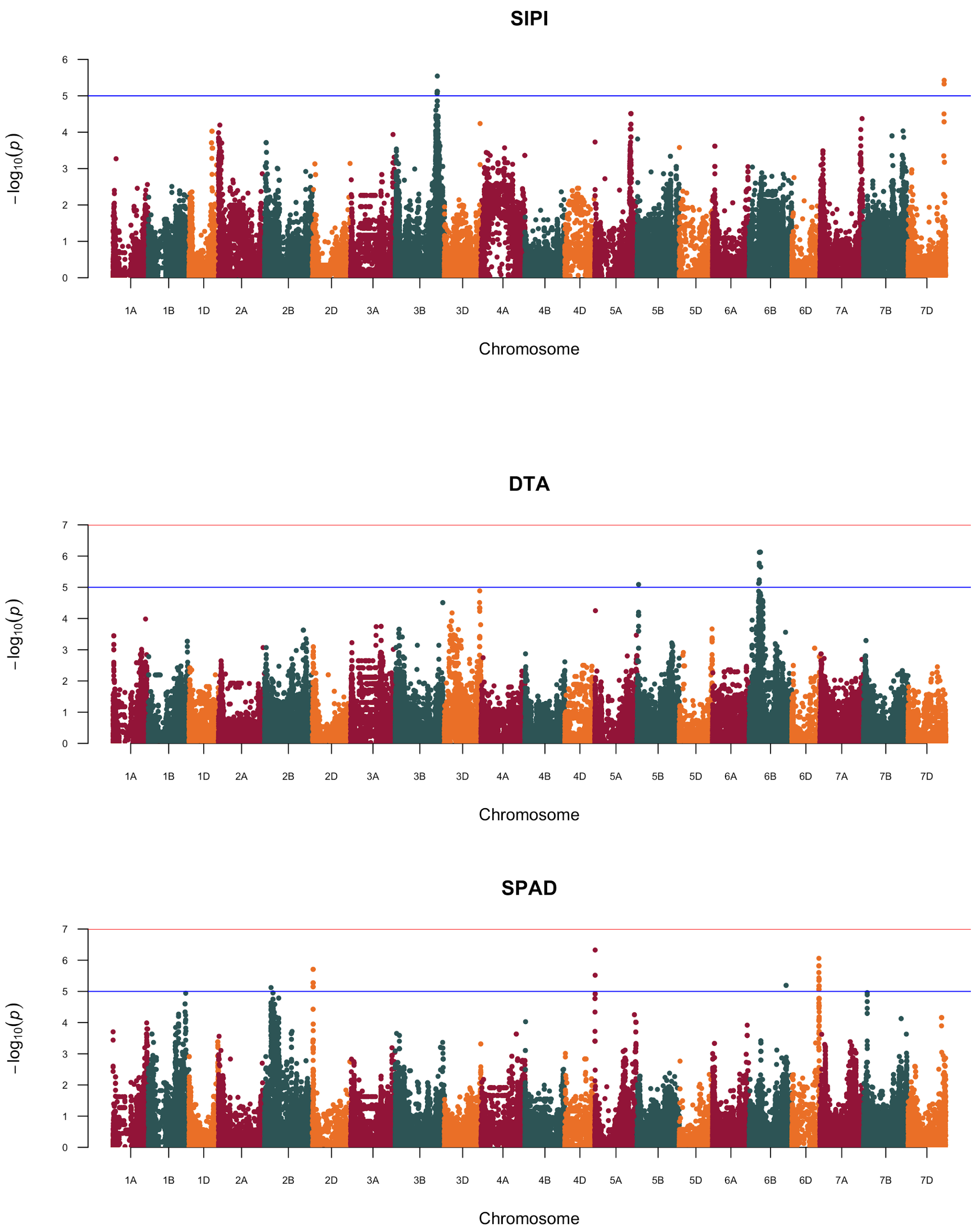
SIPI
DTA
SPAD
−log10(p)
Chromosome
1A
1B
1D
2A
2B
2D
3A
3B
3D
4A
4B
4D
5A
5B
5D
6A
6B
6D
7A
7B
7D

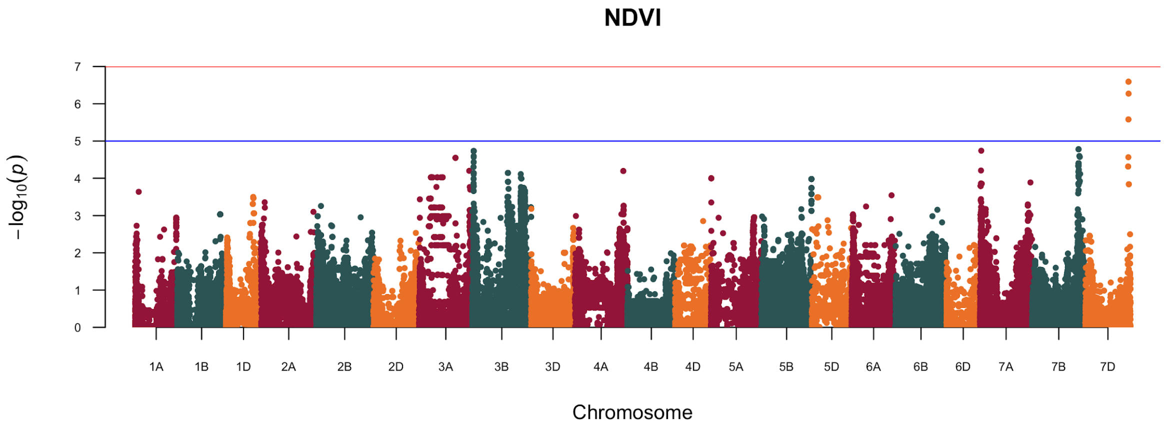
NDVI
−log10(p)
0
1
2
3
4
5
6
7
1A 1B 1D 2A 2B 2D 3A 3B 3D 4A 4B 4D 5A 5B 5D 6A 6B 6D 7A 7B 7D
Chromosome

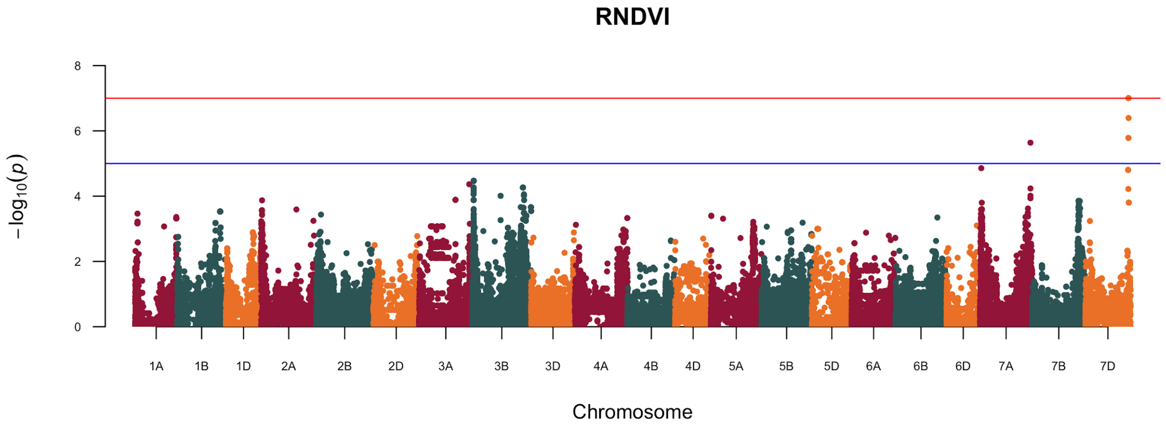
RNDVI
−log10(p)
0
2
4
6
8
1A 1B 1D 2A 2B 2D 3A 3B 3D 4A 4B 4D 5A 5B 5D 6A 6B 6D 7A 7B 7D
Chromosome

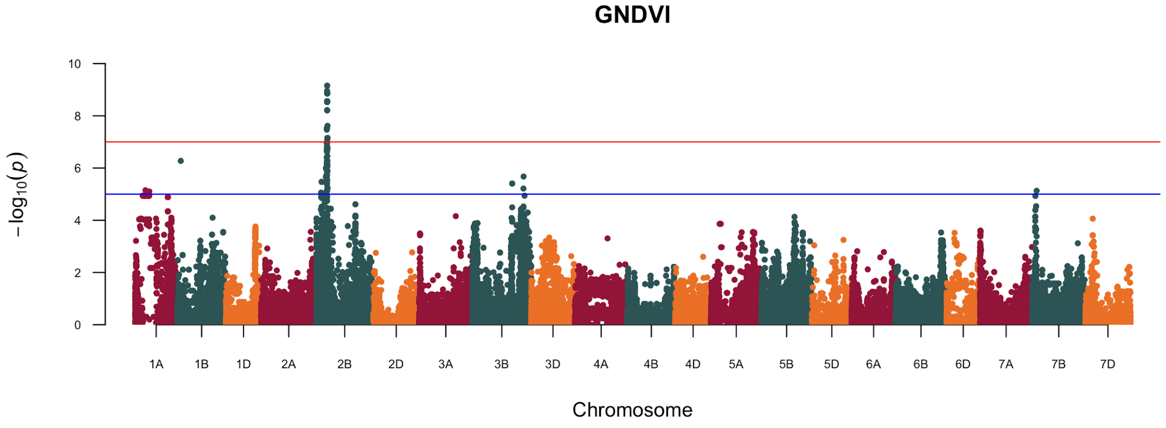
GNDVI
−log10(p)
0
2
4
6
8
10
1A 1B 1D 2A 2B 2D 3A 3B 3D 4A 4B 4D 5A 5B 5D 6A 6B 6D 7A 7B 7D
Chromosome

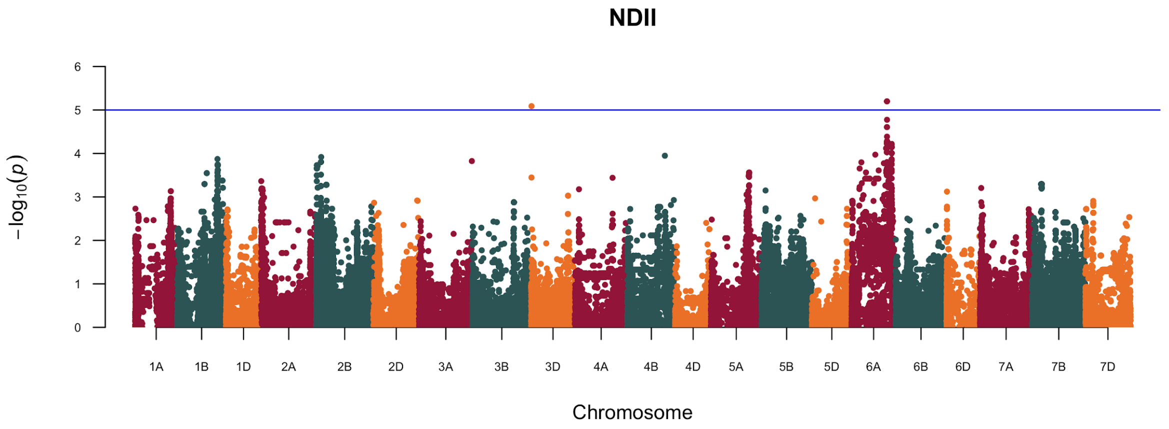
NDII
$-\log_{10}(p)$
0 1 2 3 4 5 6
1A 1B 1D 2A 2B 2D 3A 3B 3D 4A 4B 4D 5A 5B 5D 6A 6B 6D 7A 7B 7D
Chromosome

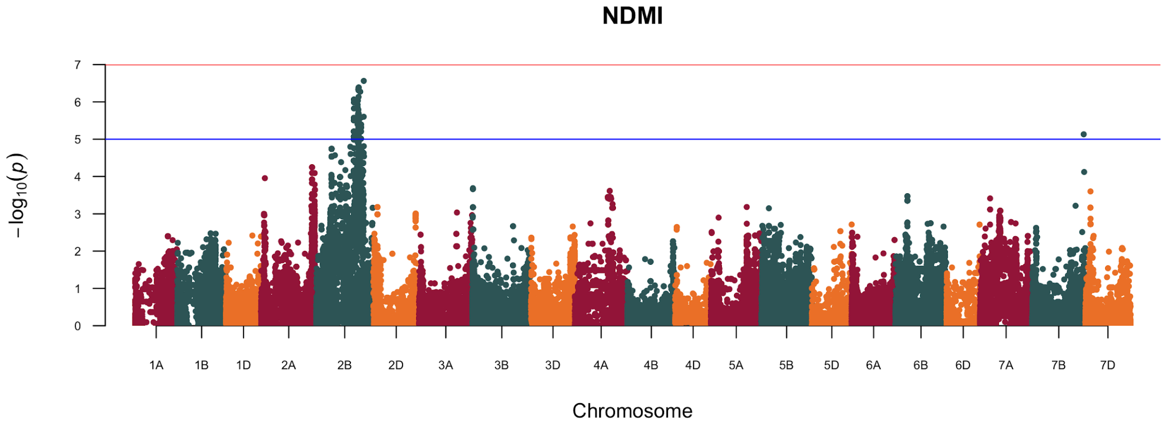
NDMI
$-\log_{10}(p)$
0 1 2 3 4 5 6 7
1A 1B 1D 2A 2B 2D 3A 3B 3D 4A 4B 4D 5A 5B 5D 6A 6B 6D 7A 7B 7D
Chromosome

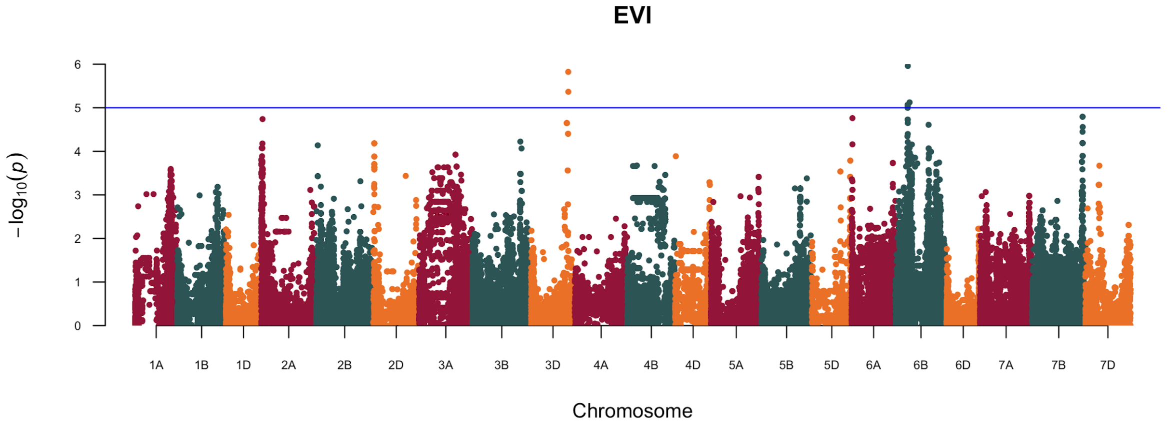
EVI
$-\log_{10}(p)$
0 1 2 3 4 5 6
1A 1B 1D 2A 2B 2D 3A 3B 3D 4A 4B 4D 5A 5B 5D 6A 6B 6D 7A 7B 7D
Chromosome

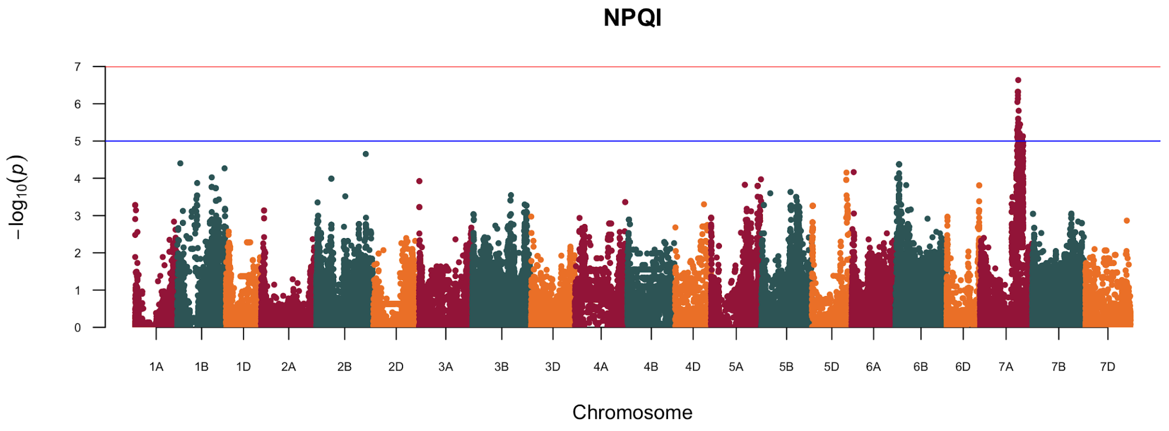
NPQI
7
6
5
4
3
2
1
0
−log10(p)
1A 1B 1D 2A 2B 2D 3A 3B 3D 4A 4B 4D 5A 5B 5D 6A 6B 6D 7A 7B 7D
Chromosome

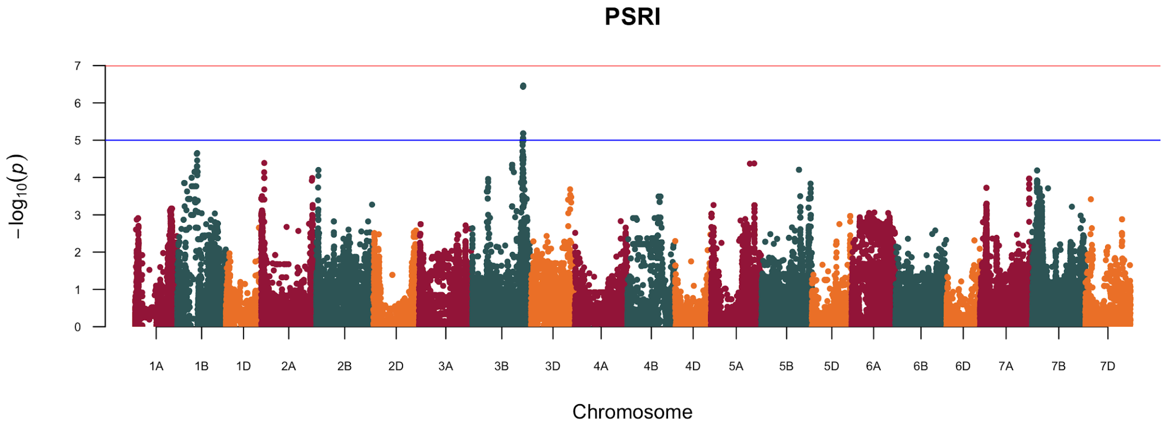
PSRI
7
6
5
4
3
2
1
0
−log10(p)
1A 1B 1D 2A 2B 2D 3A 3B 3D 4A 4B 4D 5A 5B 5D 6A 6B 6D 7A 7B 7D
Chromosome

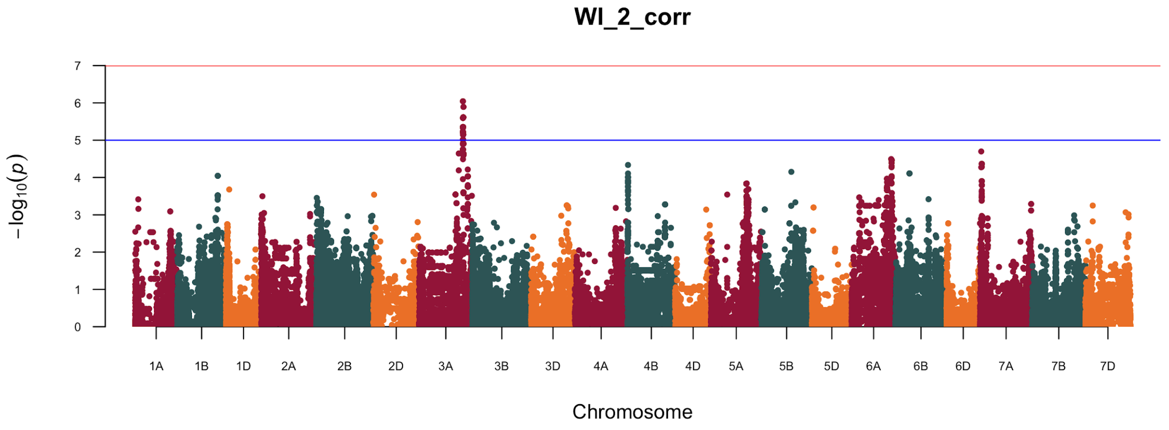
WI_2_corr
7
6
5
4
3
2
1
0
−log10(p)
1A 1B 1D 2A 2B 2D 3A 3B 3D 4A 4B 4D 5A 5B 5D 6A 6B 6D 7A 7B 7D
Chromosome

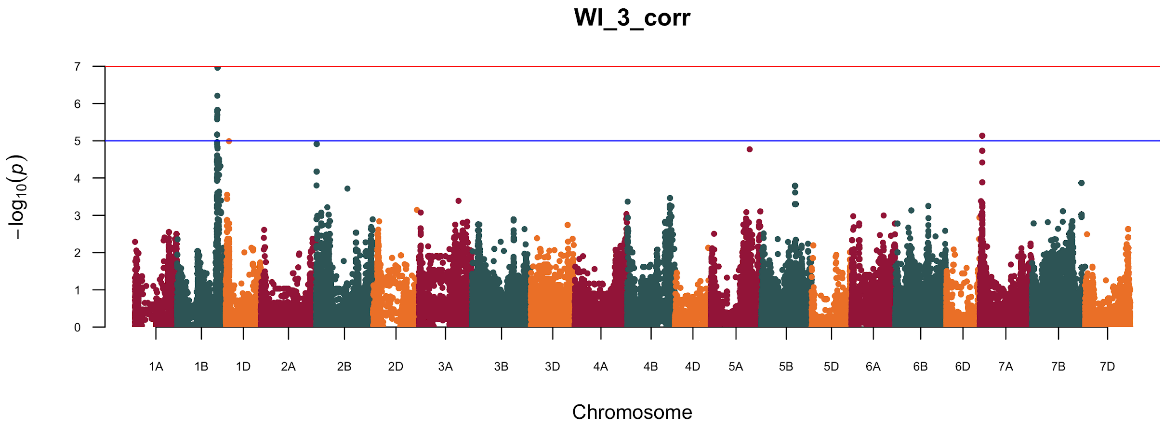
WI_3_corr
−log10(p)
0
1
2
3
4
5
6
7
1A 1B 1D 2A 2B 2D 3A 3B 3D 4A 4B 4D 5A 5B 5D 6A 6B 6D 7A 7B 7D
Chromosome

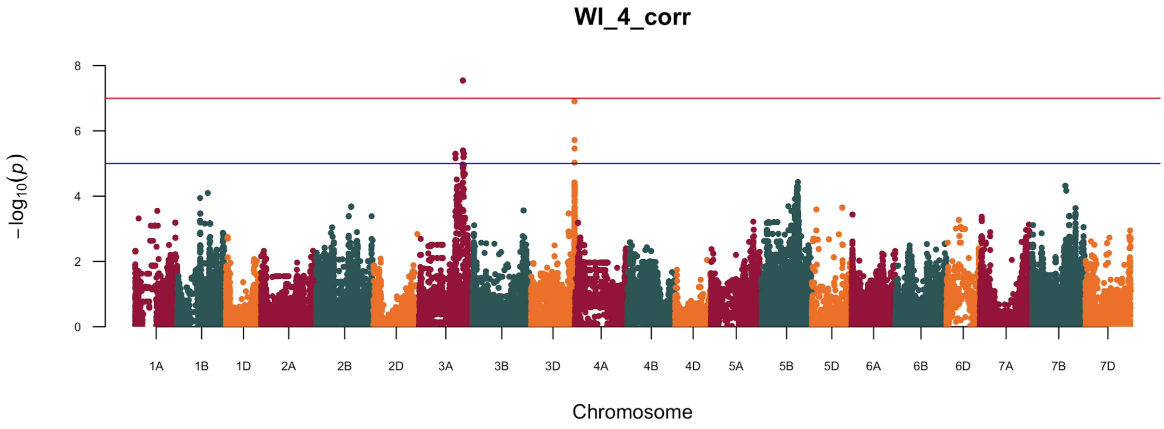
WI_4_corr
−log10(p)
0
2
4
6
8
1A 1B 1D 2A 2B 2D 3A 3B 3D 4A 4B 4D 5A 5B 5D 6A 6B 6D 7A 7B 7D
Chromosome

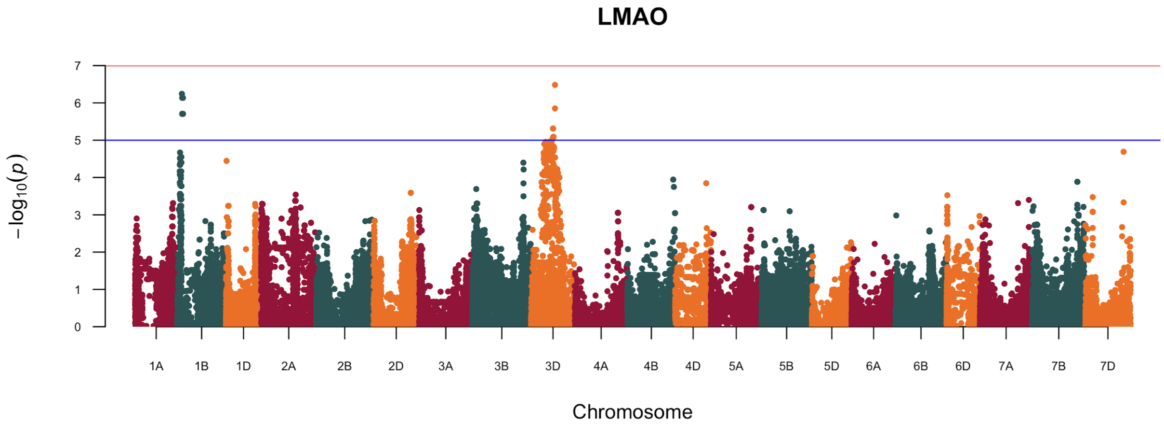
LMAO
−log10(p)
0
1
2
3
4
5
6
7
1A 1B 1D 2A 2B 2D 3A 3B 3D 4A 4B 4D 5A 5B 5D 6A 6B 6D 7A 7B 7D
Chromosome

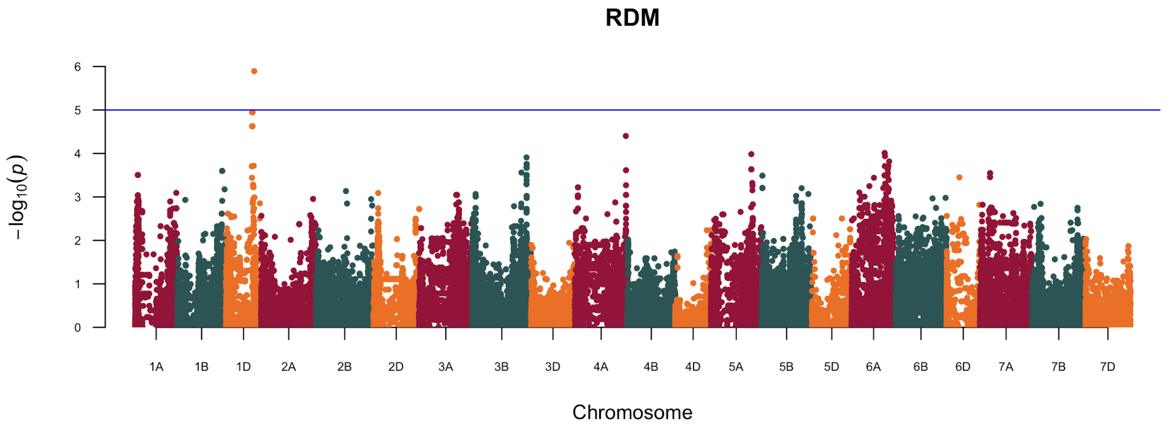

RDM
6
5
4
3
2
1
0
−log10(p)
1A
1B
1D
2A
2B
2D
3A
3B
3D
4A
4B
4D
5A
5B
5D
6A
6B
6D
7A
7B
7D
Chromosome
